# Chemical toxicity of microplastics is stronger than particle effects in *D. magna*

**DOI:** 10.64898/2026.05.12.724551

**Authors:** Simona Mondellini, Michael Schwarzer, Matthias Schott, Marvin Kiene, Bettie Cormier, Dipannita Ghosh, Martin Löder, Seema Agarwal, Martin Wagner, Christian Laforsch

**Affiliations:** Animal Ecology I and BayCEER, University of Bayreuth (UBT), Universitätsstraße 30, 95447, Bayreuth (Germany); Department of Biology, zNorwegian University of Science and Technology (NTNU), 7491 Trondheim (Norway); Macromolecular Chemistry II, University of Bayreuth (UBT), Universitätsstraße 30, 95447, Bayreuth (Germany)

**Keywords:** microplastics, extracts, *Daphnia magna*, food limitation, plastic chemicals

## Abstract

Microplastics (MP) are ubiquitous environmental contaminants with diverse physicochemical characteristics. Many studies have shown that size, shape, and polymer type are responsible for their toxicity, but this also seems to differ among MP from the same plastic type. One parameter likely contributing to these differences is plastic chemicals, a broad class of compounds intentionally or unintentionally added to plastics during their production and manufacturing. However, knowledge on the composition of plastic chemicals and their effects remains scarce. Therefore, to elucidate the chemical aspect of MP toxicity, we exposed *Daphnia magna* individuals to MP (PET, PBS, and PDLLA), cellulose, extracted particles (eMP), and methanol-based extracts of these particles for 10 days. Chemicals within such extracts were analyzed via GC-MS. This study was conducted with reduced food availability to investigate plastic effects in an environmentally relevant scenario. The introduction of a high-food control suggests that a more realistic feeding regime might exacerbate the plastic effects of the selected treatments. Our results indicated that, depending on the polymer type, plastic chemicals determine MP toxicity, which varies according to the endpoint investigated (i.e., body length, reproduction, levels of ROS and LPO). Body length, in particular, was significantly impaired by PET and PDLLA extracts, whereas reproduction was affected by most treatments. The investigated biochemical parameters (ROS and LPO) were not affected by the exposure. These results suggest that MP toxicity strongly depends on their chemical composition, whereas adverse effects due to physical properties are present independently of chemical composition across all MP types.

**Graphical Abstract:** 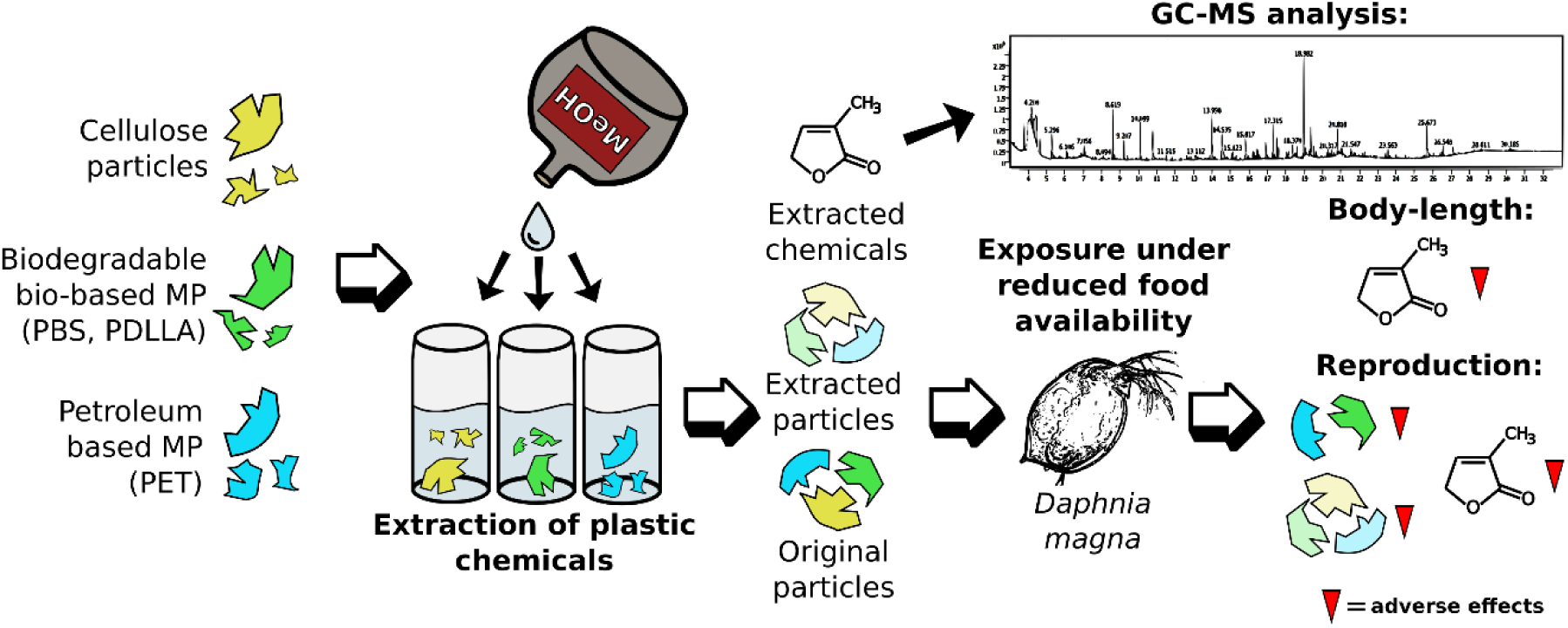

## 1. Introduction

Plastics are a diverse class of contaminants, characterized by complex chemical compositions. They are composed not only of one or more polymers but also of several additional chemicals. These chemicals can be intentionally added to enhance functionality, such as additives (e.g., fillers, flame retardants, pigments, or plasticizers) or non-intentionally added, including impurities, degradation products, or residual chemical compounds from the manufacturing process (Wagner et al. 2025). While producers are not required to disclose the chemicals used in plastic production (Beiras et al. 2021), more than 16,000 plastic chemicals have been identified in plastics (Monclús et al. 2025). At least 4,200 of these are chemicals of concern that can harm the environment or human health (Monclús et al. 2025). Some of these compounds (e.g., HBCDs, PBDEs, bisphenols, DEHP) are priority substances under the Water Framework Directive (2000/60/EC) and have also been included in the Environmental Quality Standard for surface waters (2008/105/EC; 2013/39/EU, Gunaalan, Fabbri, and Capolupo 2020).

Most plastic chemicals are not covalently bound to the polymer backbone (Zimmermann et al. 2019), increasing the likelihood that they leach from the material throughout its entire life cycle (Giametti and Finelli 2022). Given the ubiquity of plastic pollution across all environmental compartments, particularly microplastics (MP, 1–1000 µm, ISO 24187:2023) inevitably interact with biota, causing adverse effects that depend on shape, size, and polymer type (Schwarzer et al. 2022). However, the adverse effects caused by plastic chemicals may be even more severe, although only a few studies have considered them as drivers of MP toxicity (e.g., Zimmermann et al. 2020, Beiras et al. 2021, Giametti and Finelli 2022, Brehm et al. 2023). Furthermore, the number and types of plastic chemicals can vary widely across the different applications of the final plastic product (Capolupo et al. 2023). Hence, materials made of the same polymers do not necessarily have the same chemical composition (Stevens et al. 2024), making a comprehensive hazard assessment of them difficult. The high diversity in chemical composition could therefore explain the inconsistent outcomes in MP ecotoxicological studies (e.g., De Felice et al. 2019, Trotter et al. 2021, Sørensen et al. 2023). Degradation processes in the environment lead to leaching of chemicals from particles into surrounding water, where they enter the aqueous phase, potentially increasing their mobility, bioavailability, and subsequent bioaccumulation in biota compared with exposure by ingestion of MP particles containing the same compounds (Jang et al. 2021). Consequently, leached plastic chemicals may exert stronger biological effects than the particles themselves (e.g., Schrank et al. 2019, Gunaalan et al. 2020, Capolupo et al. 2023). Furthermore, smaller MP are characterized by a higher surface-to-volume ratio, which can promote faster leaching of associated chemicals (Frazzetto et al. 2022, Allan et al. 2022), leading to an interplay of degradation, fragmentation, and chemical leaching in the environment. Given the large number and high complexity of plastic chemicals, knowledge of their effects remains limited to date, even though they may substantially contribute to their overall hazard.

To address this knowledge gap, we compared the effects of MP, extracted MP (eMP, i.e., particles depleted of extractable chemicals), and extracts (i.e., containing extractable chemicals from MP particles) from the same MP on the freshwater crustacean *Daphnia magna*. *D. magna* was chosen as it is a well-established model organism with a filter-feeding behavior, which makes it highly susceptible to ingesting MP from the water column.

This study was carried out using irregular fragments (< 20 µm) of one petroleum-based (PB-) plastic: poly(ethylene terephthalate) (PET); and two biodegradable and bio-based (BB-) plastics: poly(butylene succinate) (PBS), and poly(D-lactic acid) (PDLLA). The effects of PB-and BB-plastics were compared to determine whether the latter (and their chemicals) cause effects similar to those of PET, as investigated in earlier studies (e.g., An et al. 2025, Serra et al. 2025, Oscar et al. 2025, Mondellini et al. 2026). BB-plastics are increasingly introduced into the market as “environmentally friendly” alternatives to conventional plastics (Shruti et al. 2019). However, to achieve physical properties comparable to those of the conventional plastics they are intended to replace, these so-called bioplastics require a similar, or even greater, amount of additional chemicals across all stages of production (Capolupo et al. 2023, Savva et al. 2023), making research on their chemical toxicity urgently needed. The MP effects were also compared to those of cellulose, a naturally occurring polymer in the same size range, to exclude the possibility that potential effects are elicited by particle exposure per se. Accordingly, the aim of this study was threefold: (i) to compare the effects of MP, eMP, and their extracts to identify the main component driving MP toxicity, (ii) to investigate the potential differences between the effects induced by PB- and BB-MP and those of their associated chemicals; and (iii) to analyze the effects of MP exposure under reduced food availability.

The reduced food availability scenario was chosen to better reflect environmentally relevant conditions, as standard testing conditions with *ad libitum* feeding may underestimate the hazard posed by MP and their associated chemicals to biota in natural settings by mitigating their potential effects (Silva et al. 2022, Ghosh et al. 2025, Lyu et al. 2022, Mondellini et al. 2026).

## 2. Materials and Methods

### 2.1 *Daphnia magna* cultivation

The individuals of *D. magna* used in this experiment were from a well-established in-house husbandry. The chosen clone was BL2.2, which originated from a pond (Oud Meren) in Leuven (Belgium) and has been kept in husbandry since 1997 (Imhof et al. 2017). For the experiment, animals were randomly selected from the third brood of age-synchronized mothers that were held in 1.5 L glass beakers (Weck GmbH u. Co. KG, Wehr-Öflingen, Germany). All animals were maintained in M4 artificial medium (Elendt and Bias 1990) with the addition of 4.48 µg/L of selenium dioxide (SeO_2_) and kept at 20±0.5 °C with a photoperiod of 14 h light : 9 h dark with 30 minutes of dusk and dawn. The *D. magna* culture was fed *ad libitum* with the green algae *Acutodesmus obliquus.* The mothers were separated from their offspring 24 h before the start of the experiment to prevent further hatching and to ensure the use of similarly aged neonates (<24h) in the experimental procedure.

### 2.2 Particle suspension and extraction of plastic chemicals

In this study, we tested MP made from three plastic types (fragments < 20 µm) and cellulose fragments (microcrystalline powder, 20 µm, CAS: 9004-34-6, Sigma Aldrich Chemie GmbH, Steinheim, Germany) as a natural particle control. Fragments of two BB plastics (PBS, BioPBS FZ91PM, Mitsubishi Chemicals, Japan; molecular weight (Mw) = 2.19 × 10^5^ g mol^−1^; PDLLA, NatureWorks, USA; Mw = 1.95 ×10^5^ g mol^−1^) and one PB plastic (PET, NEOGROUP, Lithuania; Mw = 9.77 ×10^4^ g mol^−1^) were produced in-house by milling (Ultra-Centrifugal Mill ZM 200, Retsch, Haan, Germany) and sieving (<20 µm, ALPINE Air Jet sieve e200 LS, FRITSCH, Idar-Oberstein, Germany). The particle size distributions of the original MP and cellulose particles (see Mondellini et al. 2026 for results) were determined via a combination of LD (laser diffraction) and DIA (dynamic image analysis) (FLOWSYNC, Microtrac MRB, Montgomeryville and York, USA).

It is essential to note that the materials used in this study were derived from pre-production pellets, which do not contain the same levels of chemicals as those eventually incorporated into final plastic products, which tend to have higher levels of intentionally and unintentionally added compounds (UNEP, 2023). We chose to use pre-production pellets rather than fragments from ground plastic materials to better compare and interpret the results with a previous study on *D. magna* using the same materials (Mondellini et al. 2026).

The extracts were prepared via solvent-based extraction (Zimmermann et al. 2020) by incubating approximately 500 MP mL^−1^ (mass equivalent based on the respective polymer density) in three 50 mL batches of methanol (LiChrosolv hypergrade for LC-MS, ≥99.9%, Supelco, Merck KGaA, Darmstadt, Germany). After 45 min of extraction in an ultrasonic bath, the glass vials containing the incubated particles were left to settle for 15 min. 100 mL of the pooled supernatant was then removed, filtered through a 0.2 µm syringe filter (Filtropur S, PES membrane, Sarstedt AG & Co. KG, Nümbrecht, Germany), and divided into two new 50 mL glass vials. The methanol was then evaporated under a gentle flow of N_2_ for approximately 6 h (Reacti-Therm III, Thermo Scientific, Waltham, Massachusetts, USA) until 1 mL of extract remained. This volume was transferred to 1.5 mL glass vials, further evaporated under N_2_ (Caliper Turbo Vap LV evaporator, Marschall Scientific, Hampton, USA), and was stopped before complete evaporation. We then added 1 mL DMSO (CAS: 67-68-5, Merck KGaA, Darmstadt, Germany) and completed the evaporation of the methanol. DMSO has a higher boiling point than methanol (189 and 64.7 °C, respectively), meaning that methanol is removed without substantial DMSO evaporation.

These 1-mL extracts contained the chemicals extracted from 50,000 MP of the respective polymer, equivalent to 138, 90, 96, and 30 mg of PBS, PLA, PET, and cellulose, respectively. Methanolic extracts represent a worst-case scenario for chemical leaching in a freshwater system (Zimmermann et al. 2020). The particles used to create the extracts (i.e., the extracted particles that should not contain other methanol-extractable compounds) were left to dry in the glass vials and then used as extracted MP (eMP) for the exposure experiments. A procedural blank was included. This procedural blank (DMSO control) was prepared following each step of the extracts’ preparation without the addition of any particle. Briefly, three batches of 50 mL of methanol (LiChrosolv hypergrade for LC-MS, ≥99.9%, Supelco, Merck KGaA, Darmstadt, Germany) were incubated for 45 min in an ultrasonic bath, then they were left to settle for 15 min. 100 mL were then removed and filtered through a 0.2 µm syringe filter (Filtropur S, PES membrane, Sarstedt AG & Co. KG, Nümbrecht, Germany), and divided into two new 50 mL glass vials, which were then evaporated first under a gentle flow of N_2_ (ca. 6 hours), then the remaining methanol was transferred in 1.5 mL glass vials to proceed with the evaporation. Before the evaporation was completed, we added 1 mL of DMSO and then completed the methanol evaporation.

Before the start of the experiment, particle suspensions were prepared by adding 100 mg of each particle to 100 mL of autoclaved M4 medium. Prior to use, stock suspensions were shaken for 48 h at 100 rpm. A haemocytometer (Neubauer Improved, Brand GmbH & Co. KG., Wertheim, Germany) was used to determine particle concentration in the stock suspensions.

### 2.3 Particle characterization

#### 2.3.1 Gas Chromatography/Mass Spectrometry for Extracts Characterization

To avoid signal interference from DMSO, GC-MS analysis was carried out on methanol extracts prepared as described above, with minor modifications: the amount of MP and the volume of methanol were reduced to one-fifth of the quantities previously used (i.e., 10 ml of methanol). After extraction (see paragraph 2.2), ∼ 6.6 mL of the methanol supernatant from each vial was collected and combined in a glass beaker to obtain a final volume of 20 mL. The 20 mL extract was filtered through a 0.2 µm syringe filter (Filtropur S, PES membrane, Sarstedt AG & Co. KG, Nümbrecht, Germany) and then divided into two new 10 mL glass vials. Methanol was removed by gentle evaporation under a nitrogen (N₂) stream (Reacti-Vap I #TS-18825 Evaporation Unit, Thermo Fisher Scientific Inc.) for approximately 6 h, until a final volume of 250 µL was reached. The final concentration of these extracts (80-fold dilution) was lower than that used during exposure (100-fold dilution); however, given the qualitative nature of the analysis, this difference is unlikely to have influenced the results.

Half a microlitre of each methanol extract was injected by an AOC20i+s autosampler (Shimadzu, Kyoto, Japan) splitless into the GC-MS (GC2030 gas chromatograph connected to a QP2020NX mass spectrometer; both Shimadzu). The GC oven was fitted with a low-polar phase; cross-linked diphenyldimethylpolysiloxane capillary column (Rtx-5MS; length = 30 m, inner diameter = 0.25 mm, film thickness = 0.25 µm; Bad Homburg v. d. Höhe, Germany) and the oven temperature was increased from 70 to 280 °C at a rate of 5 °C min^−1^, then held for 7 minutes. Helium was used as the carrier gas at a linear velocity of 50 cm s^−1^.

Mass spectrometry was measured in TIC mode in a mass range between 35 and 550 m/z. Blank measurements were performed before and after each measurement to verify the absence of carryover. A reference mixture of C7–C33 n-alkanes (Sigma-Aldrich, St. Louis, USA) was used for calculating the Kováts index of the unknown substances (Kováts 1958). All Peaks were analyzed by a NIST search (Version 2.3). If the Kováts Indices of the unknown substances and their respective 3 best NIST hits did not match, the NIST result was deemed unreliable and deleted (see Table 1). Only compounds uniquely present in one extract and absent in the others were reported in the Results section.

#### 2.3.2 Nuclear magnetic resonance

Both MP and eMP were analyzed with 1H (300 MHz) Nuclear magnetic resonance (NMR) to determine if the chemistry of the plastic material has been changed by the extraction process. Spectra were recorded on an Ultrashield-300 spectrometer (Bruker) in CDCl_3_. The residual peak of the undeuterated component in CDCl_3_ was used for calibration.

#### 2.3.3 Scanning electron microscopy analyses

The morphology and surface characteristics of both original MP and eMP particles were analyzed at a scanning electron microscope (SEM, JEOL JSM-IT500 Scanning Electron Microscope, JEOL Ltd., Japan). Particles were fixed to aluminum stubs (Ø 12 mm, Plano GmbH, Ernst-Befort-Straße 12, Wetzlar, Germany) on conductive carbon pads (G3347, Plano GmbH) and were coated with 2 nm C and 2 nm Pt (EM ACE600, Leica Microsystems GmbH, Wetzlar, Germany).

### 2.4 10-day *Daphnia magna* exposure

The exposure to 500 particles mL^−1^, or the equivalent of extracts, was carried out for 10 d. The animals were exposed in 900 mL glass beakers (Weck GmbH u. Co. KG, Wehr-Öflingen, Germany) filled with 500 mL of modified M4 medium, each containing 10 daphnids. Every treatment was replicated 5 times. The exposed animals were fed every second day with 0.5 mg C L^−1^ of the green algae *Acutodesmus obliquus.* In addition to the MP, eMP, and extracts, we included a blank control, a solvent control (DMSO, 0.01 %, 50 µL), and a high-food control, in which the organisms were fed 2 mg C L^−1^. The exposure medium, and so the respective contaminant, was exchanged every other day. After 6 d of exposure, we acquired pictures of five animals/replicate using a dissecting microscope equipped with a digital camera (M50 Leica; camera: DP26, Olympus, Hamburg, Germany; Light: KL 300 LED, Leica) to analyze morphological parameters (cellSens Dimension, v1.11, Olympus). The number of offspring produced at the first clutch was also recorded to assess reproduction alterations.

### 2.5 Biochemical analyses

Reactive oxygen species (ROS) levels were analyzed as described in Mondellini et al. (2022). In short, at the end of the 10 d exposure, organisms of each treatment were pooled and homogenized in 1 mL of potassium phosphate buffer (100 mM, pH = 7.4, with 100 mM KCl, 1 mM EDTA, 1 mM Dithiothreitol (DTT, CAS: 3483-12-3, Sigma Aldrich, Missouri, USA) and protease inhibitors 1/100 v/v (cOmplete^™^ ULTRA Tablets, Mini, *EASYpack* Protease Inhibitor Cocktail, 05892970001, Merck KGaA, Darmstadt, Germany), an aliquot of 250 µL of the homogenate was separated before centrifugation for lipid peroxidation (LPO) analyses. The remainder of the homogenate was centrifuged at 15000 g for 30 min at 4 °C, and the supernatant was used for biochemical analyses, replicated twice. Protein quantification was performed prior to analysis using the Bradford (1976) method, and the resulting protein concentration was used to normalize the data. ROS levels were quantified by assessing the fluorescence change (**λ_ex_ =** 595 nm, **λ_em_ =** 530 nm) of 2’7’ diclhorodihydrofluorescine diacetate (2’7’DCFH-DA) (Sigma-Aldrich, St. Louis, Missouri, USA), determined by the presence of pro-oxidative molecules (Deng et al. 2009). The concentration of ROS was normalized to the protein content and expressed as arbitrary units AU DCF mg proteins^−1^. LPO was assessed according to Ohkawa et al. (1979) by quantifying thiobarbituric acid reactive substances (TBARS). All the biochemical analyses were conducted on a microplate reader (Thermo Scientific Varioskan LUX 3020-82516) at λ = 595 nm for protein quantification, λ_ex_ = 485 nm and λ_em_ =530 nm for ROS quantification, and λ = 535 nm for LPO analyses.

### 2.6 Statistical analyses

All statistical analyses were conducted using R (version 4.2.2, R Core Team 2022). First, we assessed whether the particle control (cellulose), the solvent (DMSO), or the food level affected the experimental animals. Therefore, we applied linear regression models with a Gaussian distribution (Poisson distribution for offspring numbers), using the respective controls as treatment levels and the beaker as a random intercept to account for the nested design. Then, we employed pairwise comparisons after Tukey HSD with Holm adjustment for multiple comparisons, using the emmeans package (Lenth et al. 2020). We did not find significant differences between the controls, except for the higher food concentration and the number of neonates in the DMSO-treated animals. Thus, we decided to proceed with the statistical analyses using the blank and the particle control (cellulose) as the most adequate control groups.

To differentiate between the effects of plastic particles and plastic chemicals, and to compare their effects across different plastic types, we applied Gaussian-distributed linear regression models (with a Poisson distribution for offspring numbers). We used the particle type (i.e., cellulose, PBS, PDLLA, PET), particle occurrence (i.e., yes/no), and chemical occurrence (i.e., yes/no) as fixed terms with a three-way interaction and the beaker as a random intercept to account for the nested design. For instance, PBS particles containing their additives have been declared as plastic type = PBS; particle occurrence = yes; additive occurrence = yes. In contrast, the extract of PBS particles (without the particles) was declared as plastic type = PBS; particle occurrence = no; chemical occurrence = yes; and so on. Therefore, the ANOVA results were directly and easily interpretable, indicating whether particles and/or extracts affected the responses of the experimental animals. To compare results across plastic types, we used a Tukey HSD post hoc test with Holm-adjustment for multiple comparisons (Lenth et al. 2020). All models have been validated by checking the residuals for normal distribution and homogeneity of variance (Hartig, 2022). For the body length of the animals, we observed a violation of the homogeneity of the variance. Therefore, we applied a variance structure (varIdent) for the extract term using the package nlme (v. 3.1-160; Pinheiro et al. 2022). Plots were created with the package ggplot2 (Wickham et al. 2016). The significance level was set to 0.05.

## 3. Results

### 3.1 Particle characterization

SEM imaging of the particles before and after the extraction process did not reveal any visible changes in shape, size, or surface roughness in all particles except for PDLLA eMP. In the case of PDLLA, the eMP appeared to be smoother than the original MP (Figure 1; for detailed view, see Figures SI 1–4; for overview micrographs of particles; see Mondellini et al. 2026). The NMR spectra of the polymers did not show any alteration after undergoing the solvent-based extraction (Figure SI 5). GC-MS analyses revealed 19 peaks: 4 for PET, 10 for PBS, 4 for PDLLA, and 1 for cellulose (Table SI 1 & Figures SI 6–10). Of these unknown peaks, only four had Kováts indices that matched those of the three best hits from the NIST library search (Table SI 1).

**Figure 1:**
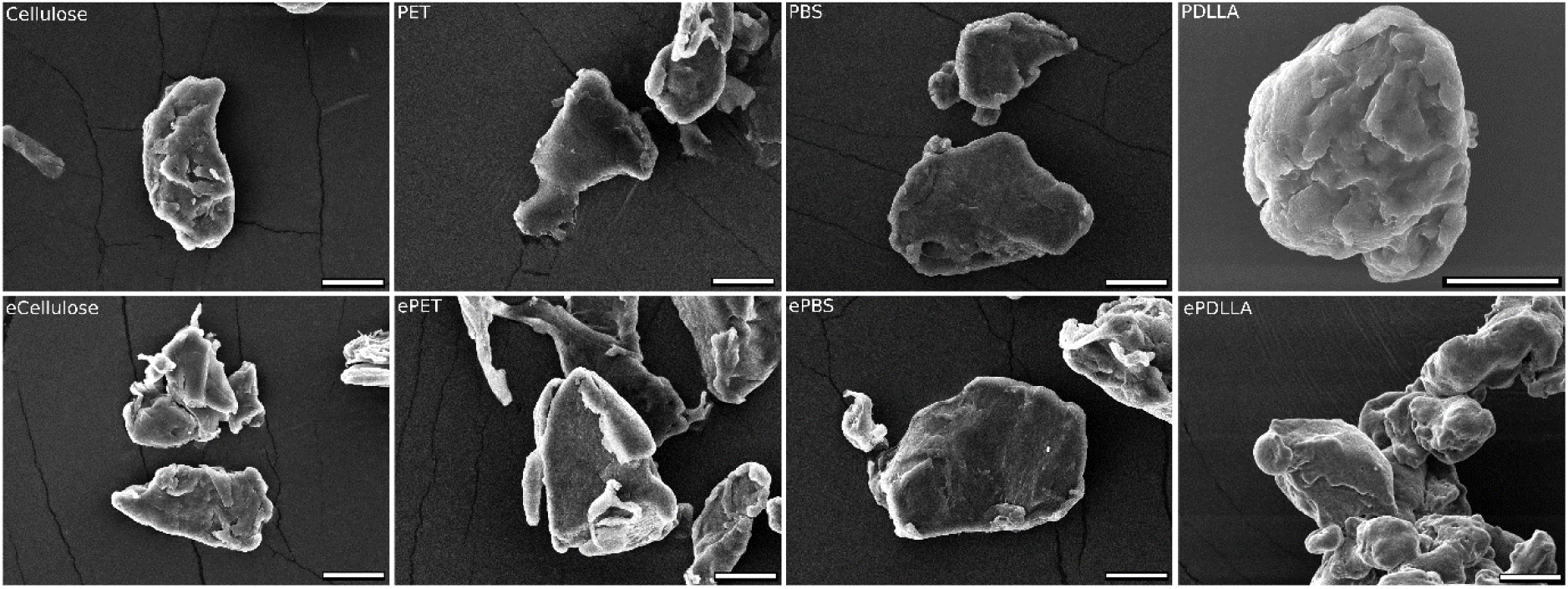
SEM micrographs of cellulose and MP particles. Top row: original particles; bottom row: particles that went through the extraction process (eMP). Scale bars = 10 µm.

### 3.2 Selection of reference control

The control comparison was conducted to assess differences among the control treatments across all selected endpoints.

Body length comparison at day 6 of all the control treatments (Figure 2) showed that animals in the food-level control were significantly larger than those in the other control treatments (cellulose-food control t = −4.687, p < 0.0001; blank-food control t = −5.381, p < 0.0001; DMSO-food control t = −7.008, p < 0.0001). The other control treatments (i.e., blank, cellulose, and DMSO) did not differ significantly from one another.

**Figure 2:**
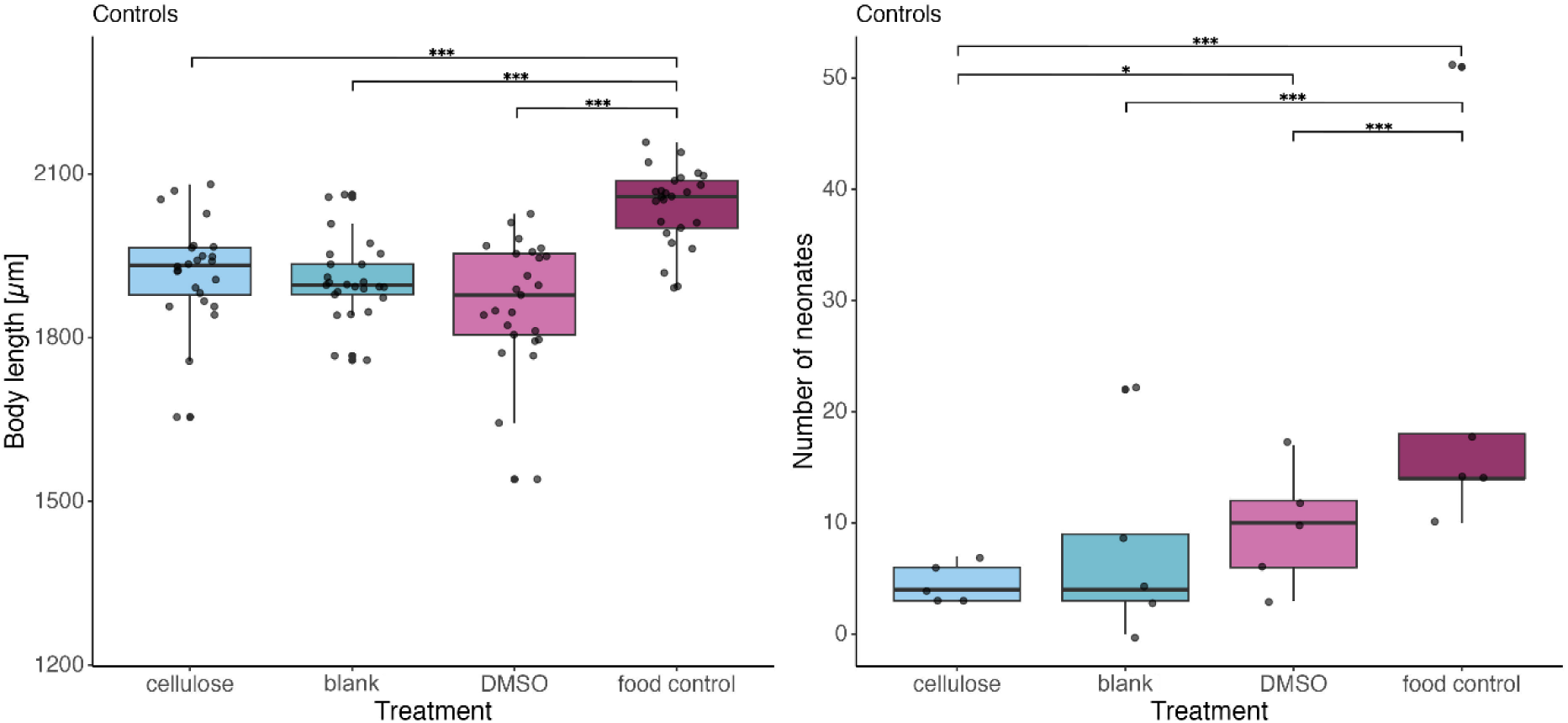
Body length (µm) measured on day 6 (N=25, with 5 individuals measured per replicate, the beaker effect was accounted for in the model) and number of hatched neonates (N=5) recorded for the control treatments, grouped by polymer and treatment type (MP, eMP, and extracts). Horizontal lines indicate the median for each treatment, while the boxes represent the interquartile range. Individual data points correspond to values obtained for each replicate. Asterisks indicate significance against the blank control (* = p ≤ 0.05, *** = p ≤ 0.001).

The comparison of reproductive success in the first brood (Figure 2) showed that animals from the food control had higher reproductive output than those from the other control treatments (cellulose-food control t = −6.689, p < 0.0001; blank-food control t = −5.482, p < 0.0001; DMSO-food control t = −4.614, p < 0.0001). Animals from the DMSO control also had higher reproductive output than those from the cellulose treatment (p = 0.01). The remaining control treatments did not differ significantly from one another.

Biochemical analyses of the control treatments showed no significant difference in ROS levels among treatments, despite the food control having lower ROS levels overall (Figure SI 11).

When considering the LPO, animals of the food control treatment had significantly higher levels of LPO, compared to animals from the other control treatments (cellulose-food control t = −6.639, p < 0.0001; blank-food control t = −6.658, p < 0.0001; DMSO-food control t =−6.745, p < 0.0001; Figure SI 11).

Because control treatments did not differ significantly across all analyzed endpoints except the food control, we compared all plastic treatments to cellulose (the natural particle control, referred to as the within-treatment comparison) and to the blank control (referred to as the within-polymer comparison).

### 3.3 Body length at day 6

Investigations on the body length after exposure (Figure 3) showed significant effects determined by particle type, type of extract, and the interactions of these parameters with themselves and the treatment (all p < 0.0001; further details in the SI Table 2-4).

**Figure 3:**
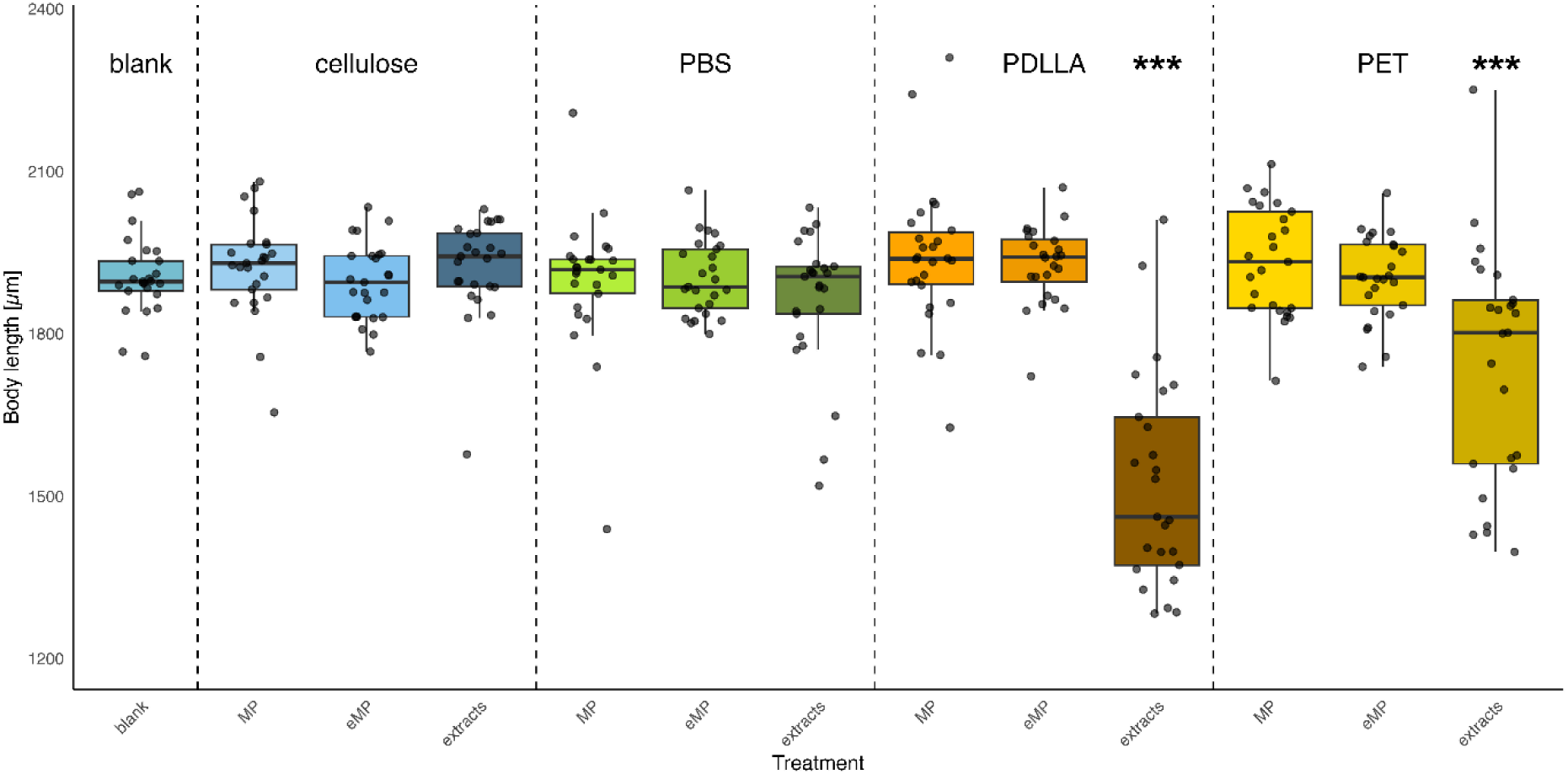
Body length (µm) measured on day 6, grouped by polymer and treatment type (MP, eMP, and extracts). Horizontal lines indicate the median for each treatment, while the boxes represent the interquartile range. Vertical dashed lines separate the different polymers. N=25, with 5 individuals measured per replicate, the beaker effect was accounted for in the model. Individual data points correspond to values obtained for each replicate. Asterisks indicate significance against the blank control (*** = p ≤ 0.001).

#### 3.3.1 Within treatment

The comparison of the different polymers within the same treatment type revealed significant differences only for the extracts. In particular, animals from the PDLLA and PET treatment (PDLLA: t = 12.37, p < 0.0001; PET: t = 5.62, p < 0.0001) were significantly smaller than those exposed to cellulose and PBS (PBS-PDLLA t = 10.49, p < 0.0001; PBS-PET t = 3.73, p = 0.0004). Comparing PDLLA and PET, PDLLA animals were smaller (t = −6.75, p < 0.0001).

#### 3.3.2 Within polymer

Pairwise comparisons within each polymer showed significant effects only for PET and PDLLA. In both cases, organisms exposed to the extracts were significantly smaller than the blank control (PDLLA: t = 11.81, p < 0.0001, D = 3.3; PET: t = 5.06, p < 0.0001), than those exposed to the respective eMP (PDLLA t = 12.54, p < 0.0001; PET t = 5.00, p < 0.0001) and MP (PDLLA t = −12.92, p < 0.0001; PET t = −5.91, p < 0.0001).

### 3.4 Reproductive success at the first brood

Investigation of the reproductive success of the first brood after exposure (Figure 4) highlighted significant negative effects of the treatment, particle type, and the interactions of treatment and particle type, treatment and extracts, particle type and extracts, and the three-way interaction (Table SI 2-4).

**Figure 4:**
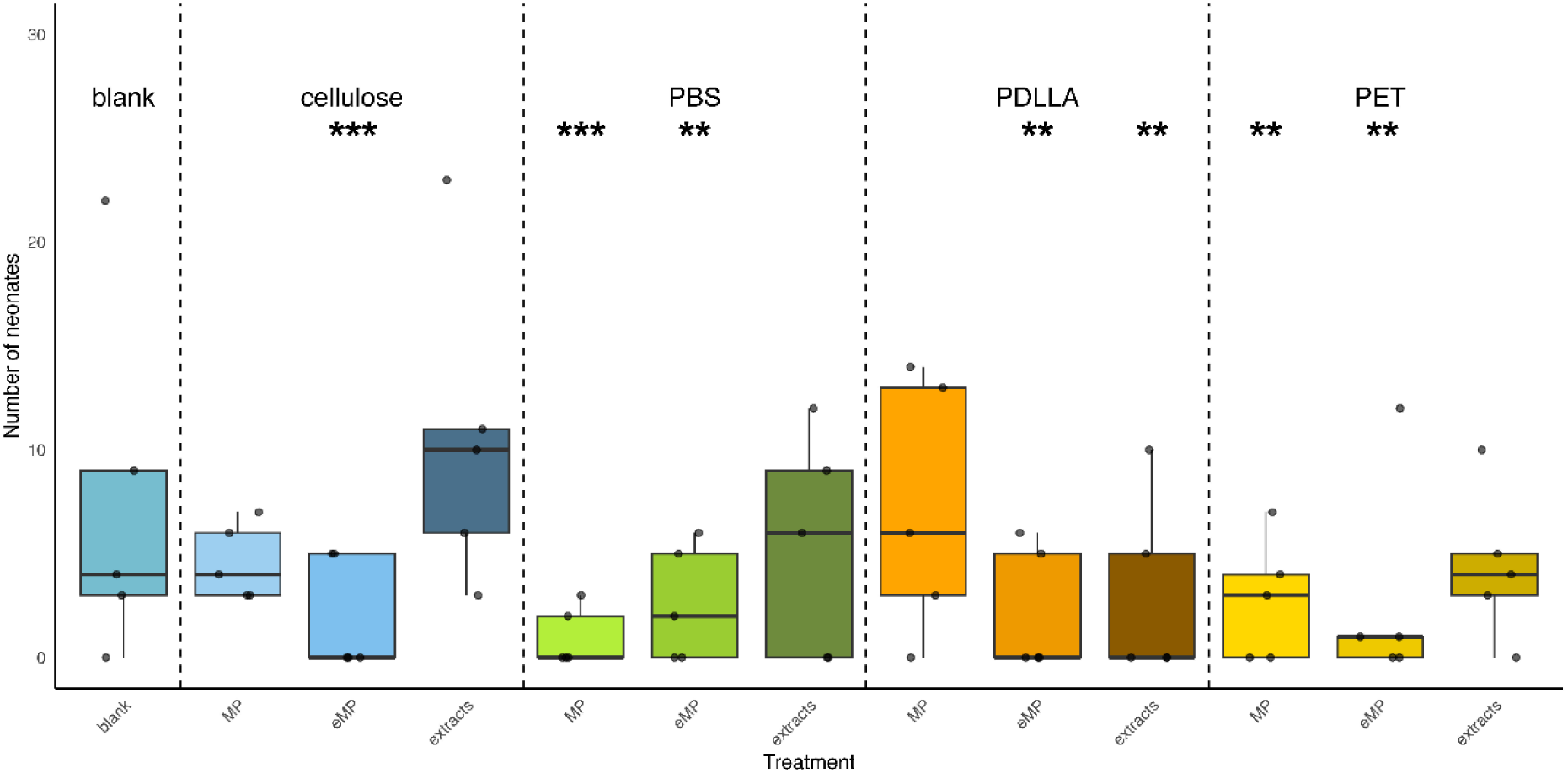
Number of neonates produced in each treatment beaker (N = 5) at the first clutch, grouped by polymer and treatment type (MP, eMP, and extracts). Horizontal lines indicate the median for each treatment, while the boxes represent the interquartile range. Individual data points correspond to values obtained for each replicate. Asterisks indicate significance against the blank control (** = p ≤ 0.01; *** = p ≤ 0.001).

#### 3.4.1 Within treatment

The reproductive outcome of the first brood of the different treatment types (Figure 4) was significantly different within the MP and extract treatments, but not within the eMP treatments. Among MP treatments, organisms exposed to PBS MP produced significantly fewer offspring than those exposed to cellulose (p = 0.0009). In contrast, PDLLA MP showed a higher reproductive output than those exposed to PBS (p = 0.0002) and PET (p = 0.01). Among the extract treatments, all polymers produced fewer neonates compared to cellulose (PBS: p = 0.01; PDLLA: p = 0.0001; PET: p = 0.002).

#### 3.4.2 Within polymer

Comparison within each polymer type revealed several significant differences (Table SI 2-4). Considering PBS, both MP (p = 0.0001) and eMP (p = 0.003) produced fewer neonates than the blank control. Additionally, PBS MP produced fewer neonates than the extracts (p = 0.002). A similar pattern was observed for PET, in which both MP and eMP treatments produced fewer neonates than the blank control (p = 0.008).

For PDLLA, the pattern was less consistent. The blank treatment showed higher reproductive output than the extracts (p = 0.002) and eMP treatments (p = 0.009). Furthermore, the PDLLA MP treatment produced a higher reproductive output than the extracts (p = 0.01) and the eMP treatment (p = 0.003).

For cellulose, the eMP treatment produced fewer neonates compared to both the blank (p = 0.0009) and the extract (p < 0.0001). In contrast, the extract treatment produced more neonates than the particle treatment (p = 0.003).

### 3.5 Biochemical analyses

The analyses of ROS levels after exposure highlighted a significant effect of the treatment (χ^2^ _3,79_ = 8.47, p = 0.037). However, no significant differences were observed in the pairwise comparisons among the treatments and polymers. When considering the levels of LPO, our analyses reported a significant effect of the treatment (χ^2^ _3,79_ = 10.306, p = 0.016) and of the particle type (χ^2^ _3,79_ = 4.075, p = 0.043). Here, pairwise comparisons between the treatments and polymers showed no significant differences (Figure SI 12–13, Table SI 2–4).

## 4. Discussion

This study aimed to elucidate the drivers of MP toxicity in *D. magna* and to compare PB- and BB-plastics by examining the original MP, eMP, and their extracts. The experimental work was conducted in a setup with environmentally realistic low food availability. Our results suggest that the severity of MP toxicity strongly depends on their chemical composition, whereas toxicity due to physical properties is present independently of chemical composition across all particles.

Specifically, body length was affected only by PET and PDLLA extracts, whereas offspring number was negatively affected by PDLLA extracts, by PET and PBS MP, and by the extracted particles (eMP). These findings indicate that the effects of PET and PDLLA are mainly driven by chemical toxicity, whereas PBS toxicity is primarily associated with the particle properties. The comparison between PB- and BB-MP revealed that both types adversely affected *D. magna*, consistent with outcomes reported for the same materials (MP particles) in a previous investigation (i.e., Mondellini et al. 2026). Notably, PBS produced the strongest adverse effects on survival and reproduction in that study, a pattern also observed here. In the present study, the adverse effects of PBS on reproduction were attributable to properties of the MP and eMP particles rather than to their chemical extracts. In contrast, PDLLA and PET primarily induced adverse effects through their extracts rather than through other particle properties. Overall, these results suggest that the tested PDLLA and PET particles exhibit greater chemical toxicity, whereas the selected PBS fragments exert stronger toxicity, potentially induced by their physical properties.

Similar to our experiment, Mondellini et al. (2026) renewed the exposure medium every other day during the exposure period, reducing potential leaching of chemicals from the particles. This may explain why PBS exerted the strongest effects in their study, as its toxicity appears to be primarily driven by physical rather than chemical factors. Consequently, the chemical toxicity of PDLLA and PET was measured in the present study. Nevertheless, we also observed physical toxicity of PET (MP and eMP) and PDLLA (eMP) in the reproduction data, emphasizing that MP toxicity arises from an interplay of physical and chemical factors (Zimmermann et al. 2020, Schwarzer et al. 2025).

To further investigate the contribution of chemical toxicity, the plastic chemicals were characterized using GC-MS. The employed extraction protocol was designed to recover most extractable chemicals from the original material using an organic solvent (i.e., methanol; Zimmermann et al. 2020). As GC-MS is limited to volatile compounds, further studies should therefore employ non-target high-resolution LC-MS to detect non-volatile compounds in plastic extracts. Although all identified substances are typically described as non-toxic or of low toxicity (i.e., Fuoco and Finne-Wistrand 2020, ECHA 2018,2021a and 2021b, NCBI, Sigma Aldrich, 2026; see SI for further details), no data are available on their potential hazard in the context of MP and environmental samples, where they appear as mixtures with uncertain combined toxicity. Plastics constitute a complex class of synthetic pollutants, and many of the toxic effects reported in the scientific literature may be attributable to plastic chemicals rather than to plastic particles per se (Gao et al. 2025). Although relatively few studies have investigated chemical extracts from plastics, several reports have documented the effects of MP leachates (e.g., Schiavo et al. 2020, Qiu et al. 2023, Jang et al. 2024).

While the effects of MP on *Daphnia* reproduction and body length have been widely reported in the scientific literature (e.g., Schwarzer et al. 2022, Brehm et al. 2023, Funke et al. 2024), the negative effects of eMP, including from cellulose, on reproduction in our study were unexpected. One possible reason for this observation could be the change of surface characteristics caused by the extraction process. However, qualitative SEM particle characterization analyses showed that eMP did not undergo visible surface alterations before and after the extraction process, except for PDLLA. SEM characterization revealed that the particles before and after extraction maintained similar size, shape, and surface roughness, except for PDLLA, whose eMP appeared to have a smoother surface than the original MP. Therefore, SEM analyses did not support the assumption that the biological responses of organisms exposed to either MP or eMP were attributable to changes in the surface morphology of these particles. In the case of PDLLA, however, this may be different. MP surface characteristics, such as surface roughness, could influence their potential toxicity (Burrows et al. 2020). For example, rough and sharp particle edges may damage cellular membranes, leading to cytolysis and necrosis (Choi et al. 2020). In addition, rougher particle surfaces have been associated with enhanced physical interaction and inflammatory responses, whereas smoother particles may be taken up by organisms more frequently than rougher ones (Khan and Jia 2023, Kim et al. 2021). Furthermore, it cannot be excluded that the extraction process also altered additional particle properties, such as surface chemistry, charge, or hydrophobicity, that were not measured in this study and that may have affected the exposure outcome (Lambert et al. 2017).

In this study, we hypothesized that all cellulose treatments could be considered controls, but the cellulose eMP treatment adversely affected reproduction, whereas the original cellulose particles did not. Therefore, the extraction process most likely affected the outcomes in this study. In fact, when considering the cellulose treatment exclusively, it is possible to observe that, while cellulose eMP reduced reproduction, the extracts increased it. This suggests that a compound present in the cellulose extracts may be responsible for the observed increase in reproduction. During the extraction process, cellulose eMP may have been stripped of this compound, thereby revealing an underlying physical toxicity that remained masked in the original cellulose particles, which had not undergone extraction. It should be noted that the extracts were dissolved in DMSO, which showed a slight stimulatory effect on the number of neonates compared to the blank control. Consequently, part of the difference between cellulose eMP and extracts may be attributed to the presence of DMSO in the latter.

However, the negative effect of cellulose eMP relative to the blank control remains more difficult to explain. One possible explanation could be the presence of trace residues or impurities in the methanol used in the extraction process. However, this explanation appears unlikely, as the solvent used was of very high purity (LiChrosolv®, Supelco, hypergrade for LC-MS, ≥99.9% measured by GC). Furthermore, the EC_50_ and the NOEC (no observed effect concentration) reported for methanol in *D. magna* are 18.26 mg L^−1^ and 122 mg L^−1^, respectively (ECHA 2020). In our study, assuming that all particles absorbed 5% of methanol and that all particles were 20 µm in size, the maximum resulting methanol concentration would have been 104.72 µL L^−1^, a level that should not have elicited adverse effects on the exposed organisms (in reality, most particles were ≤20 µm, leading to an even lower hypothetical concentration). Therefore, the observed adverse effects of cellulose and plastic eMP treatments are most likely attributable to the physical toxicity of these particles, although changes in particle composition or surface properties induced by the extraction process itself cannot be ruled out.

Contrary to our expectations, the investigated biochemical parameters, ROS and LPO levels, were not affected by any of the plastic treatments. MP are known to cause oxidative stress in several organisms (e.g., Liu et al. 2019, Silva et al. 2021, Roy et al. 2023, Oduro et al. 2025). The stable ROS levels, together with the lack of lipid-related oxidative damage observed in our study, might be attributable to the tested concentration, which could be too low to elicit an alteration in the daphnids’ oxidative status (e.g., De Felice et al. 2022, Esterhuizen et al. 2024). Previous investigations that report an alteration of this endpoint often use higher MP concentrations compared to the one used in the current study (e.g., Zhu et al. 2022, Jeyavani et al. 2022). It is important to note that the concentration tested (500 MP mL^−1^) is already elevated compared to those currently found in the environment (Gupta et al. 2022), representing a possible worst-case scenario. Another possible explanation for the lack of significant ROS and LPO responses is the reallocation of energy toward detoxification and antioxidant defense systems. Such compensatory mechanisms may prevent measurable oxidative damage while diverting energetic resources away from growth and reproduction (De Coen and Janssen 2003, Barata et al. 2005), which aligns with the observed results regarding body length and number of offspring in our study. Further studies investigating the levels of key antioxidants and detoxifying enzymes should be conducted to shed light on this aspect.

To investigate the impact of low food availability, a high-food control was included in our experiment. In line with our initial expectations, animals from this control treatment grew larger and had more offspring than those in the other control treatments (i.e., blank, DMSO, and cellulose). Regarding ROS and LPO levels, we observed no significant differences in ROS, although pro-oxidant levels appeared lower in the food control than in the others. However, we recorded a significant increase in lipid damage (LPO), which could be attributed to higher lipid content in animals in the food control treatment, as they had more resources available to store as lipid droplets. Animals from all the other treatments (limited food availability) probably depleted their fatty acid reservoirs (Yang et al. 2021), thereby lowering their pool of peroxidable substrate. These differences among the control groups suggest that the limited food provided in this study may have exacerbated the negative effects of our particle treatments (Mondellini et al. 2026) and that food availability should be accounted for in future studies on the matter of MP toxicity in *D. magna* (Ghosh et al. 2025).

## 5. Conclusion

This study compared the effects of MP, eMP, and extracts from PET (PB-MP), PBS, and PDLLA (BB-MP), and cellulose on *D. magna* under food-limited conditions to enhance environmental relevance. The study had multiple aims: (i) to investigate whether the plastic chemicals (extracts), the original MP, or just the particle presence (eMP) are responsible for MP toxicity, (ii) to assess whether BB-MP are less harmful than PB-MP, and (iii) to evaluate these drivers of MP toxicity under environmentally relevant conditions. The results highlighted the critical role of associated chemicals in plastic toxicity, indicating that, for certain endpoints, the observed toxic effects can be attributed primarily to these chemical components rather than to the particles themselves. Our findings demonstrated that BB-MP cannot be considered a safer alternative to PB-MP, as they exert comparable negative effects on daphnids. Furthermore, our results underscore the importance of incorporating environmentally realistic scenarios in future assessments, such as reduced food availability, which exacerbate plastic-related effects that could otherwise remain overlooked under standardized testing protocols. However, this study also revealed some limitations that should be addressed in future studies, as discussed in the SI (see “Limitations and recommendations”). Taken together, these findings call for a reassessment of current hazard evaluation frameworks, emphasizing the need to account for chemical co-exposure and environmental realism in assessing the ecological safety of MP derived from both conventional and biodegradable bio-based plastics.

## Authors contribution statement

**Simona Mondellini*:** Conceptualization, Methodology, Investigation, Data curation, Writing - Original Draft, Writing - Review & Editing, Formal analysis, Visualization, Project administration, Validation

**Michael Schwarzer*:** Conceptualization, Methodology, Investigation, Data curation, Writing - Original Draft, Writing - Review & Editing, Formal analysis, Visualization, Project administration, Validation

**\*** contributed equally

**Matthias Schott:** Methodology, Investigation, Data curation, Formal analysis, Writing - Review & Editing

**Marvin Kiene:** Writing - Review & Editing, Formal analysis, Visualization

**Bettie Cormier:** Conceptualization, Investigation, Writing - Review & Editing

**Dipannita Ghosh:** Investigation, Writing - Review & Editing

**Martin G.J. Löder:** Investigation, Writing - Review & Editing, Validation, Project administration, Supervision

**Seema Agarwal:** Project administration, Funding acquisition, Resources, Writing - Review & Editing

**Martin Wagner:** Conceptualization, Writing - Review & Editing, Validation, Project administration, Supervision, Funding acquisition

**Christian Laforsch:** Conceptualization, Resources, Writing - Review & Editing, Validation, Project administration, Supervision, Funding acquisition

## Acknowledgements

The SEM was funded by the Deutsche Forschungsgemeinschaft (DFG GZ: INST 91/427–1 FUGG). We thank Prof. Dr. Johannes Stökl and Dr. Jacqueline Sahm from the department of Evolutionary Animal Ecology for technical support with GC-MS sample preparation, and Jana Karola Koch for helping with the analysis.

