## Supplementary Information for "Chemical toxicity of microplastics is stronger than particle effects in *D. magna*"

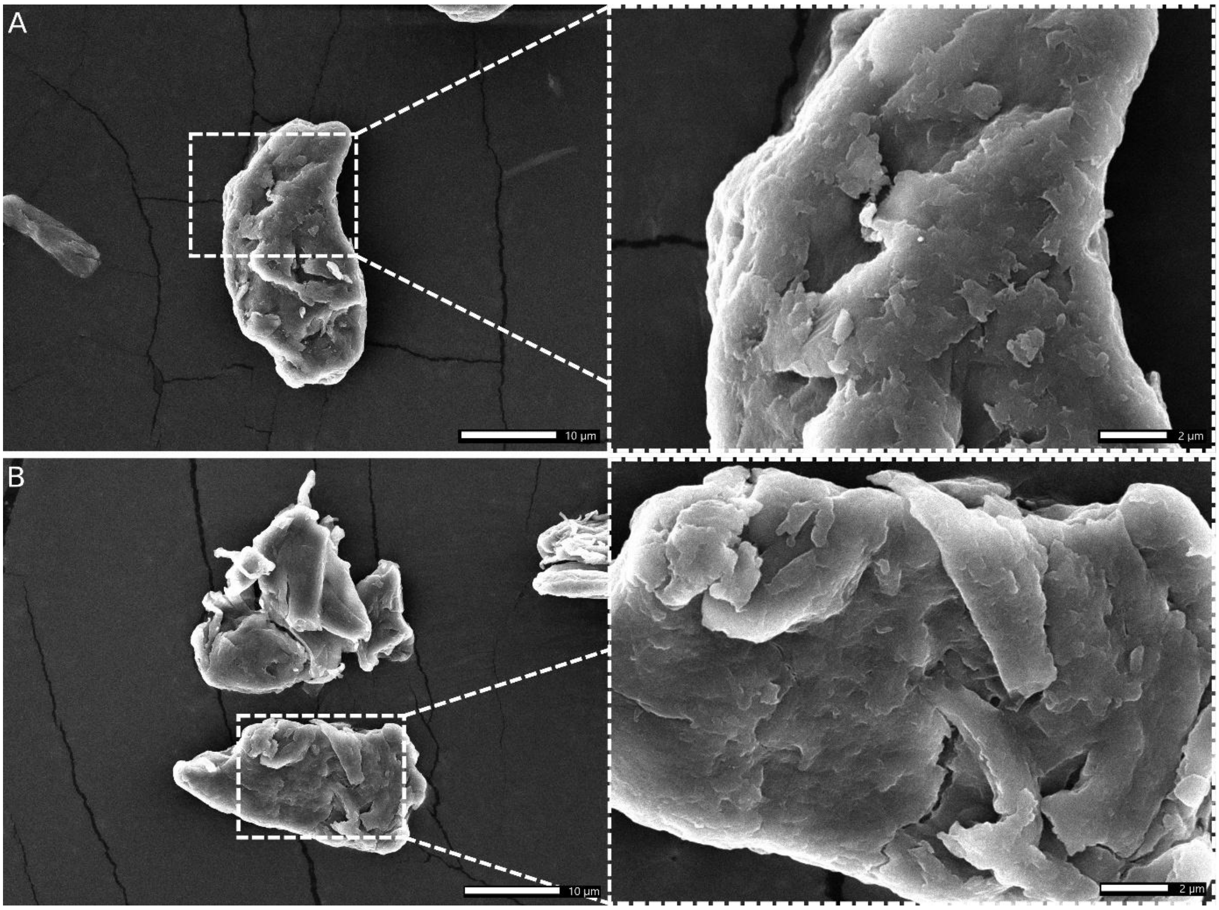


***Figure SI 1:*** *SEM images of cellulose particles. (A) pristine cellulose. (B) eCellulose. Left: Overview, scale bar = 10 µm; Right: Detail, scale bar = 2 µm.*

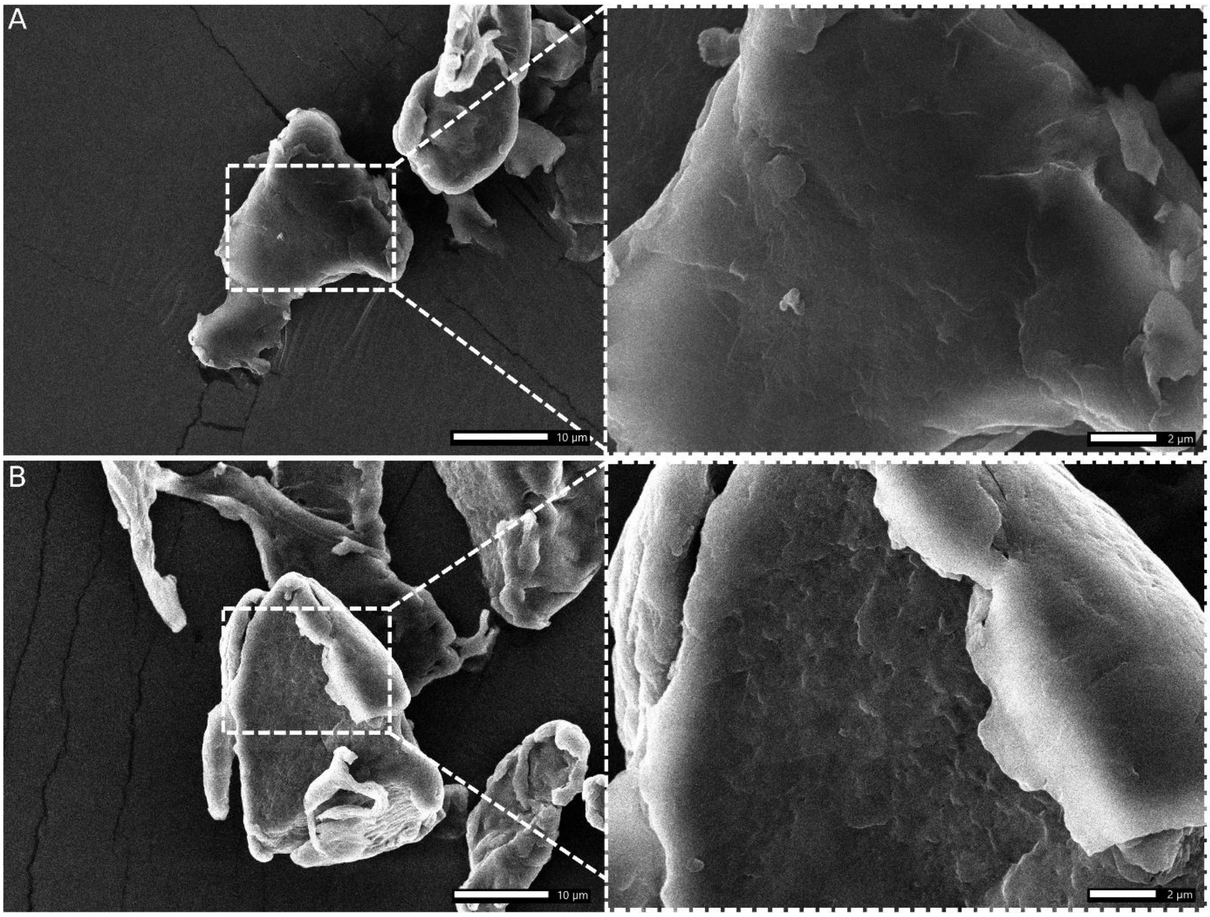


***Figure SI 2:*** *SEM images of PET particles. (A) pristine PET. (B) ePET. Left: Overview, scale bar = 10 µm; Right: Detail, scale bar = 2 µm.*

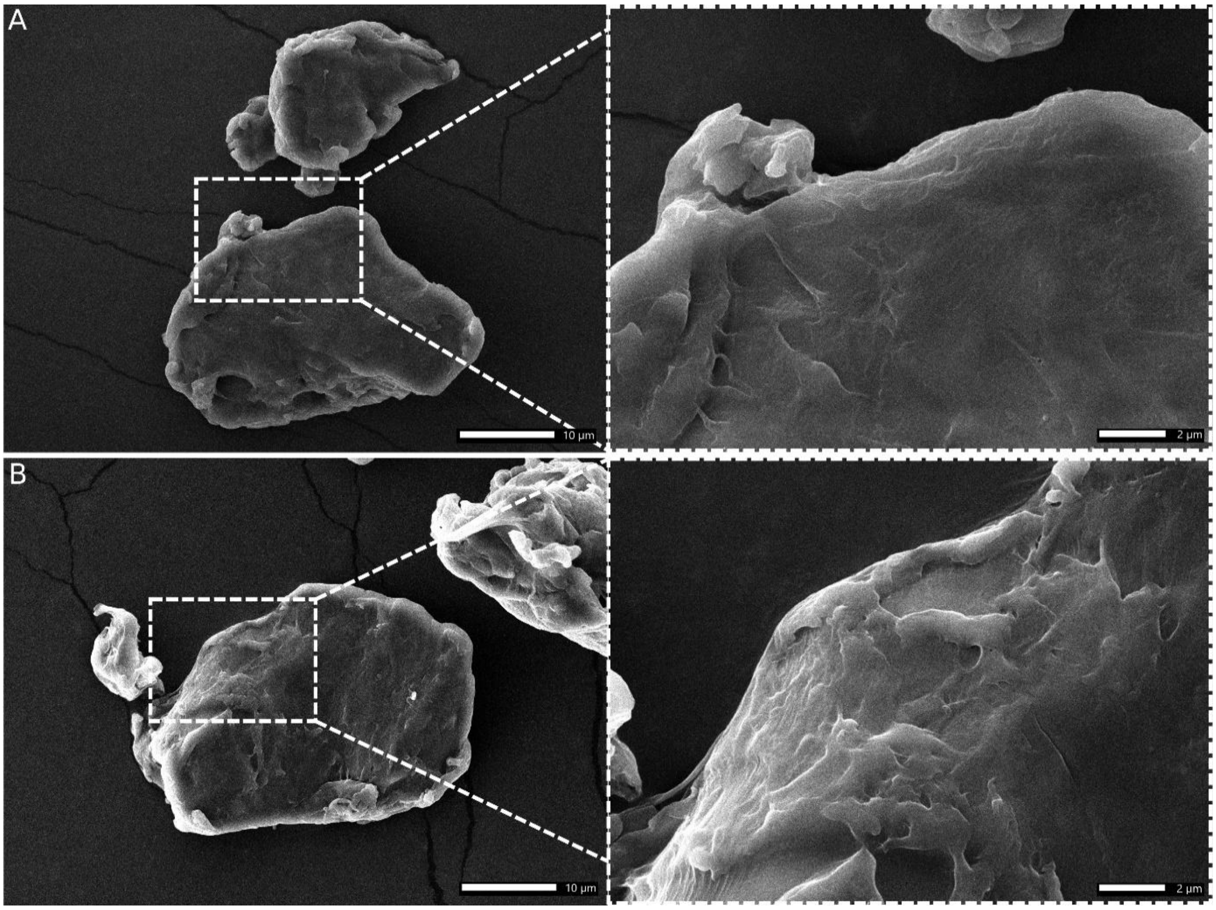


***Figure SI 3:*** *SEM images of PBS particles. (A) pristine PBS. (B) ePBS. Left: Overview, scale bar = 10 µm; Right: Detail, scale bar = 2 µm.*

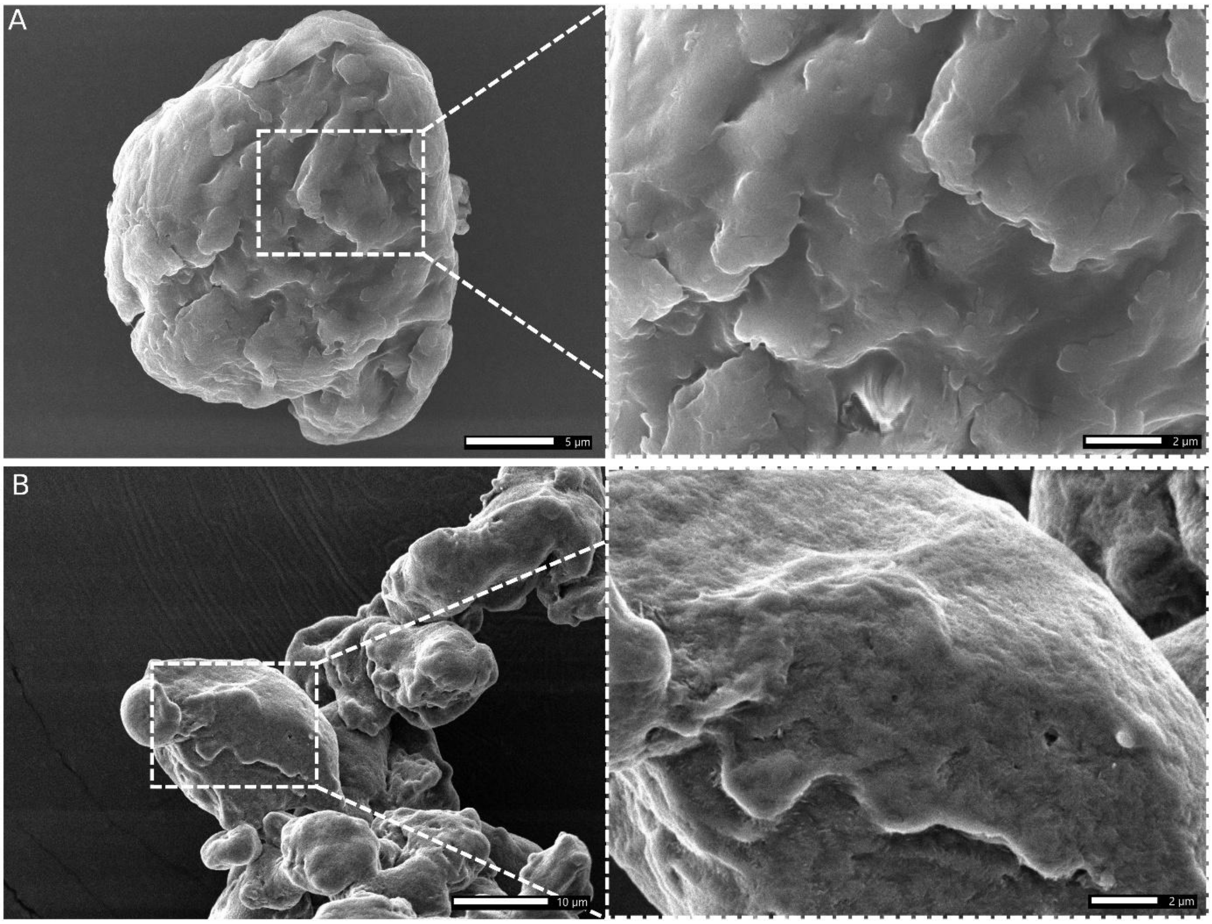


***Figure SI 4:*** *SEM images of PDLLA particles. (A) pristine PDLLA, scale bar = 5 µm. (B) ePDLLA, scale bar = 10 µm. Left: Overview; Right: Detail, scale bar = 2 µm.*

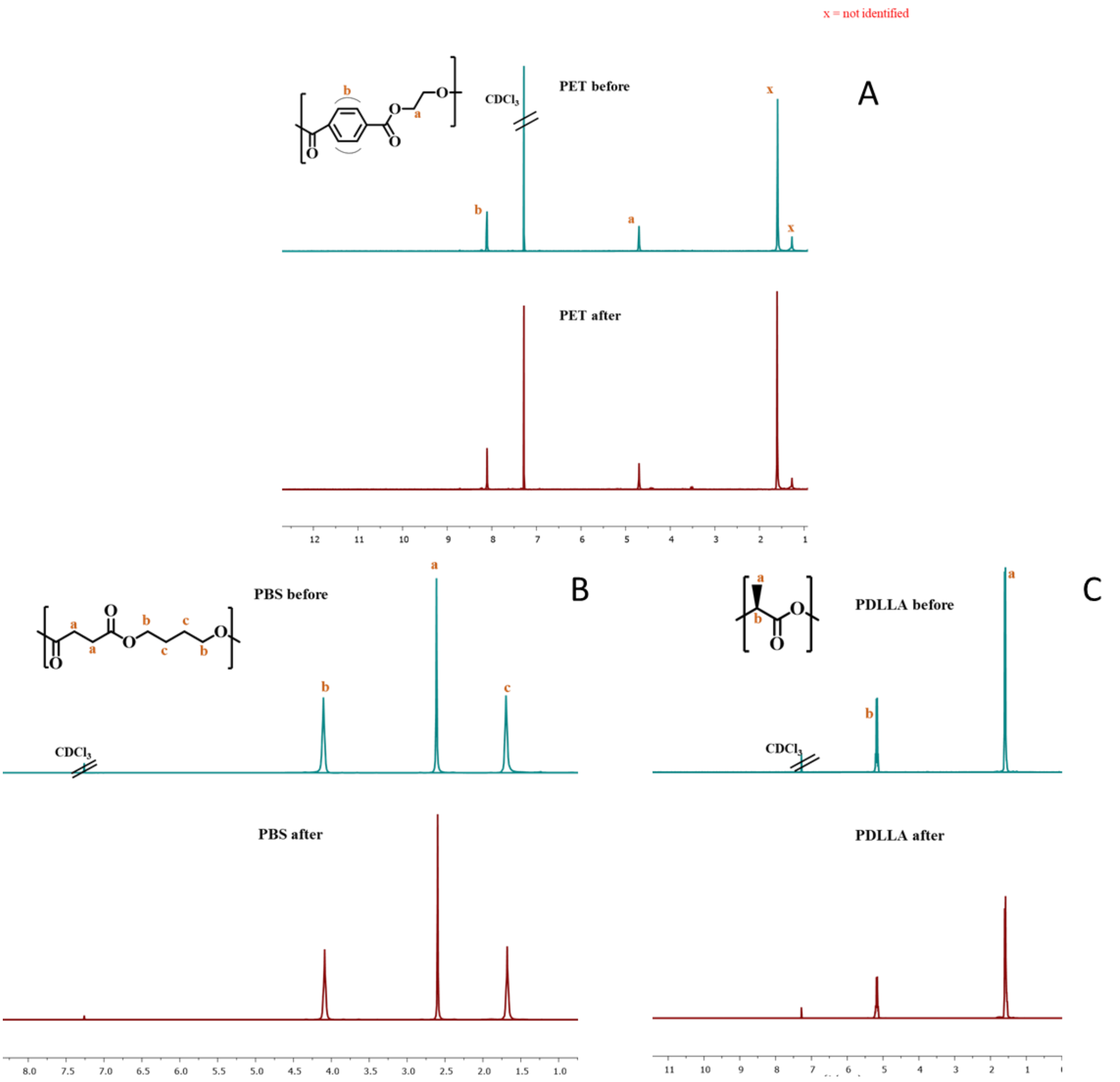


***Figure SI 5:*** *NMR spectra in CDCl₃ before (blue spectra) and after (red spectra) the methanol extraction process of A) PET, B) PBS, and C) PDLLA. Not identified peaks are indicated with x.*

**Table SI 1**: Results of the GC-MS analyses conducted on the MP extracts. AC: associated chemical, RT: retention time, m/z: mass to charge ratio of base peak, Calculated Kováts: calculated Kováts index of measured peak (Kováts 1958). S/N: signal-to-noise ratio. For corresponding chromatograms, see Fig. SI 7.

| *Polymer* | *AC* | *CAS* | *RT* | *S/N* | *m/z* | *Calculated Kováts* | *Literature Kováts* |
|---|---|---|---|---|---|---|---|
| PET | Thiodiglycol | 111-48-8 | 9.98 | 167.72 | 61 | 1184.73 | 1190.1 |
| | Peak 1 | NA | 14.76 | 109.63 | 163 | 1507.34 | 1550.4 |
| | Docosane | 629-97-0 | 21.90 | 15.28 | 57 | 2198.55 | 2200 |
| | Peak 2 | NA | 26.11 | 3.95 | 55 | 2731.44 | NA |
| PBS | Peak 3 | NA | 4.73 | 505.45 | 42 | 947.17 | NA |
| | Butanedioic acid, dimethyl ester | 123-25-1 | 6.99 | 341.49 | 115 | 1042.44 | 1035.5 |
| | Thiodiglycol | 111-48-8 | 9.97 | 98.88 | 61 | 1184.37 | 1190.1 |
| | Peak 4 | NA | 12.65 | 129.10 | 101 | 1322.50 | NA |
| | Peak 5 | NA | 15.51 | 204.13 | 115 | 1570.83 | 1705 |
| | Peak 6 | NA | 21.19 | 238.26 | 73 | 2120.00 | NA |
| | Peak 7 | NA | 21.75 | 132.37 | 115 | 2124.35 | NA |
| | Peak 8 | NA | 22.28 | 9.74 | 43 | 2243.05 | NA |
| | Peak 9 | NA | 24.98 | 397.33 | 55 | 2578.70 | NA |
| | Peak 10 | NA | 26.10 | 139.63 | 115 | 2730.92 | NA |
| PDLLA | 2(5H)-Furanone, 3-methyl- | 22122-36-7 | 5.70 | 10.50 | 41 | 988.18 | 982 |
| | Peak 11 | NA | 8.95 | 740.25 | 45 | 1134.11 | NA |
| | Peak 12 | NA | 9.00 | 378.11 | 56 | 1142.23 | NA |
| | Peak 13 | NA | 14.40 | 96.91 | 45 | 1480.53 | NA |
| *Cellulose* | Peak 14 | NA | 25.44 | 9.73 | 43 | 2641.50 | NA |

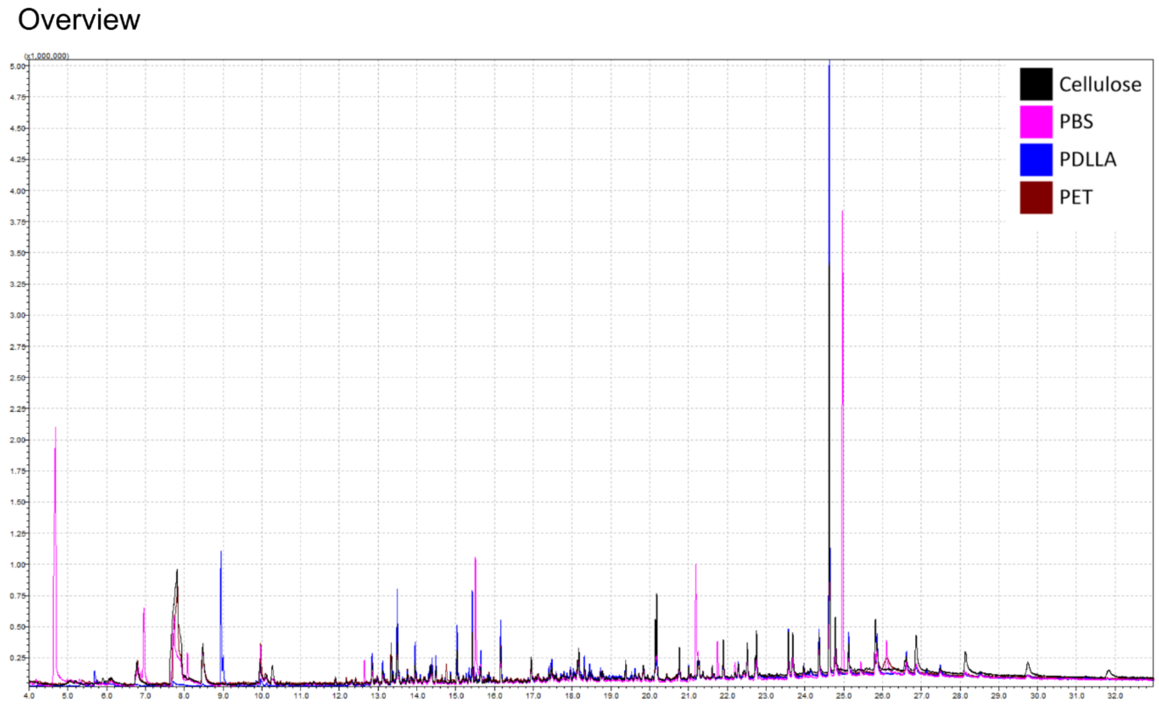


***Figure SI 6:*** *Total Ion Chromatograms of a GC-MS analysis of a sample of each treatment.*

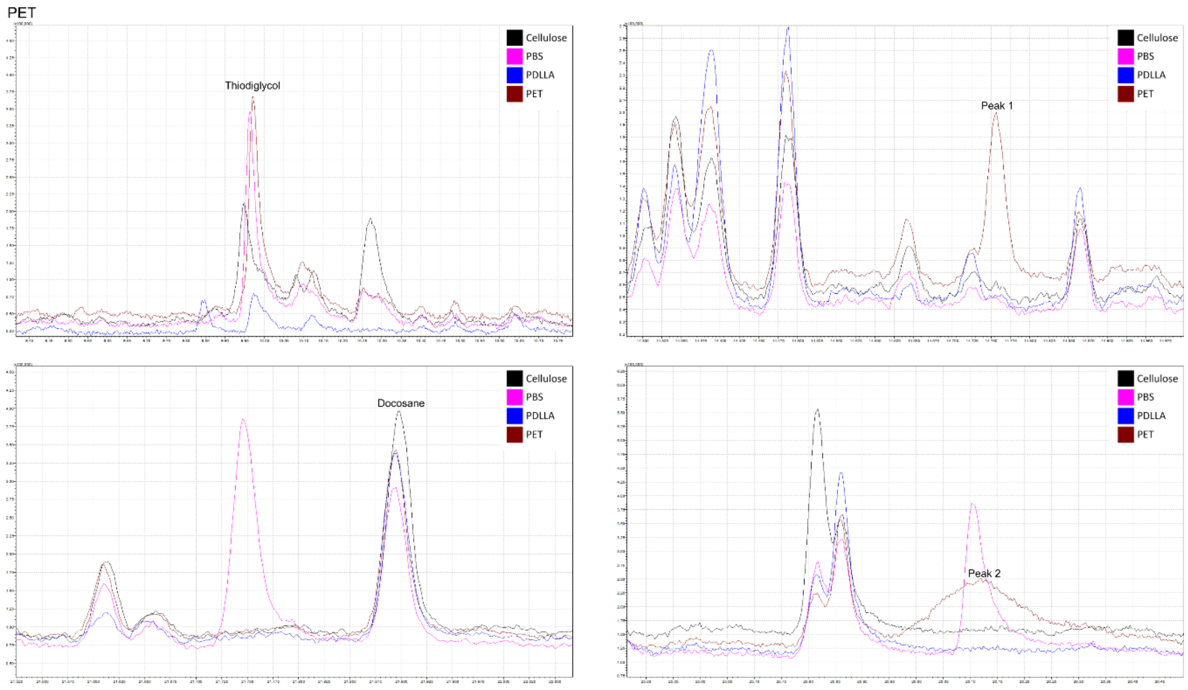


***Figure SI 7:*** *Zoom in of total Ion Chromatograms to unique peaks of the PET treatment. All listed peaks were present in all PET replicates. Peak annotations correspond to Table SI 1.*

PBS

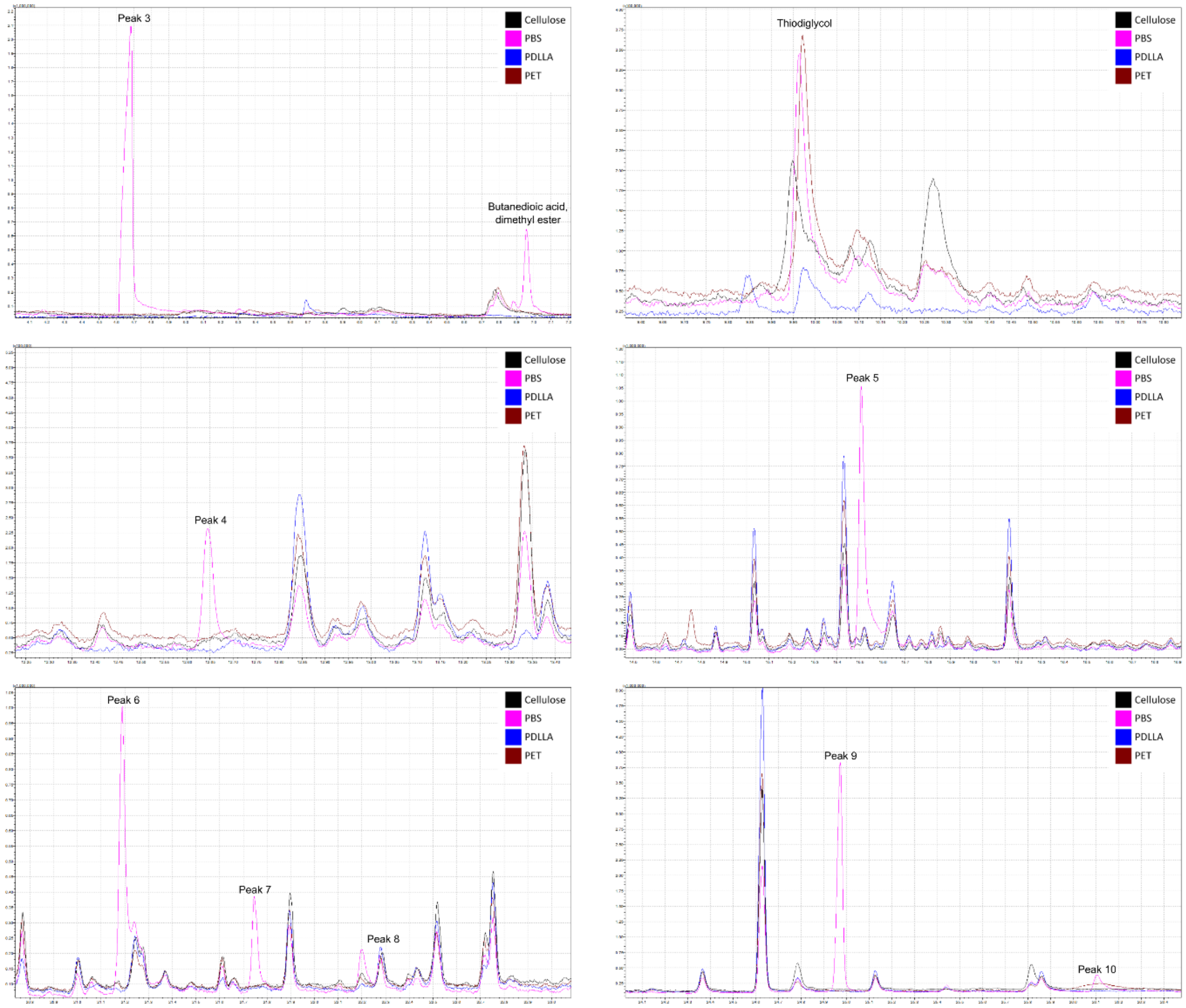


***Figure SI 8:*** *Zoom in of the total Ion Chromatograms with unique peaks of the PBS treatment. All listed peaks were present in all PBS replicates. Peak annotations correspond to Table SI 1.*

PDLLA

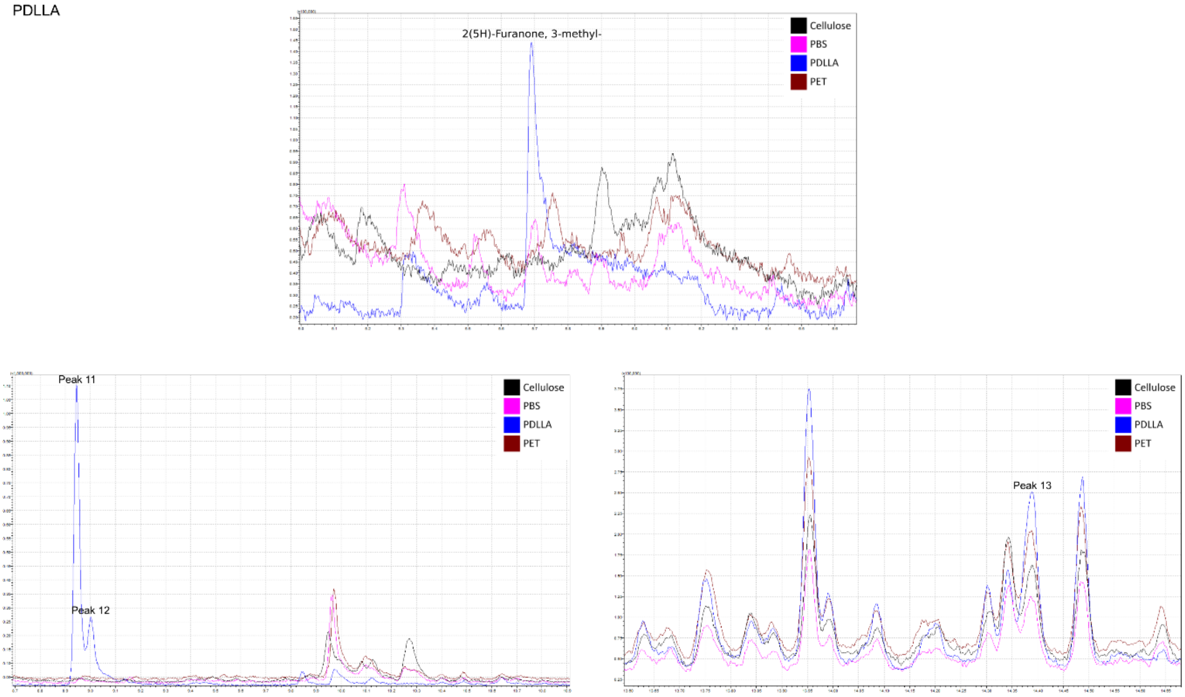


*Figure SI 9:* *Zoom in of the total Ion Chromatograms unique peaks of the PDLLA treatment. All listed peaks were present in all PDLLA replicates. Peak annotations correspond to Table SI 1.*

Cellulose

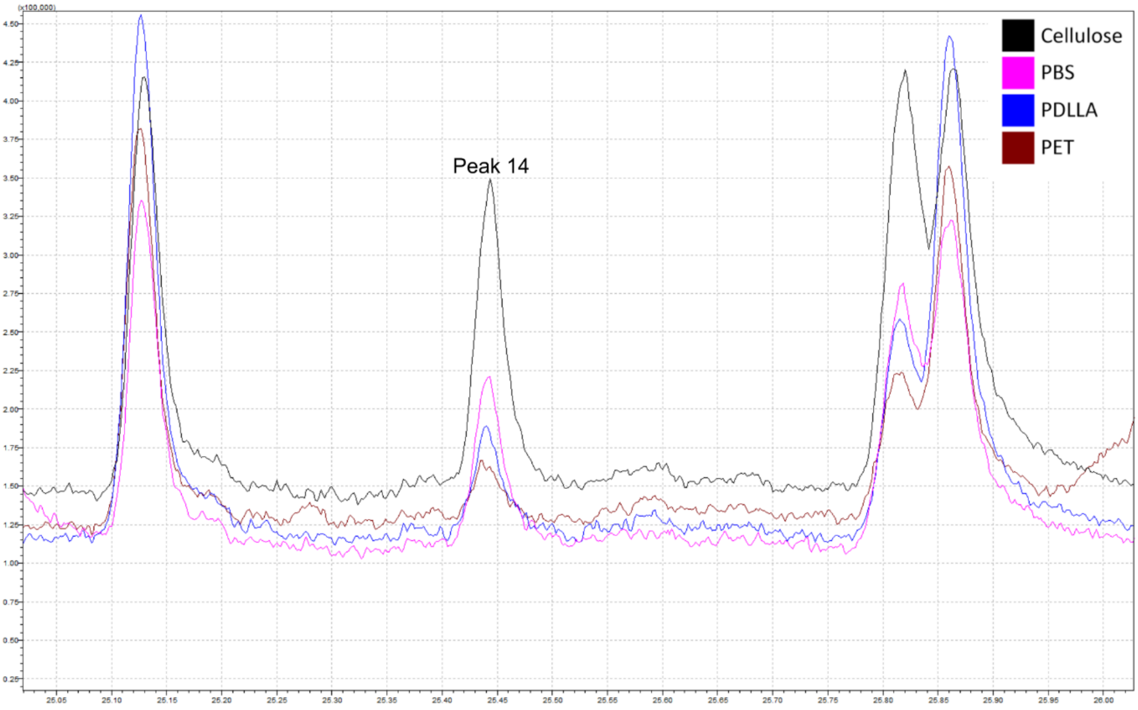

***Figure SI 10:*** *Zoom in of the total Ion Chromatograms of the unique peak of the Cellulose treatment. The peal was present in all cellulose replicates. Peak annotations correspond to Table SI 1.*

**Table SI 2:** parameters from the linear regression analyses of *D. magna'*s life history parameters obtained during the 10-day exposure. Bold values represent significance ($p < 0.05$).

| ***Response*** | ***Fixed effects*** | ***DF*** | ***$\chi^2$ /F value*** | ***p-value*** |
|---|---|---|---|---|
| ***Body length (day 6)*** | Treatment | 3 | 4.07 | 0.253 |
| | Particle type | 1 | 13.63 | **0.0002** |
| | Extracts | 1 | 30.6 | **<0.0001** |
| | Treatment:particle type | 3 | 22.0 | **<0.0001** |
| | Treatment:extracts | 3 | 45.5 | **<0.0001** |
| | Particle type:extracts | 1 | 48.2 | **<0.0001** |
| | Treatment:particle type:extracts | 3 | 44.7 | **<0.0001** |
| ***Num. of offspring*** | Treatment | 3 | 9.8 | **0.019** |
| | Particle type | 1 | 52.9 | **<0.0001** |
| | Extracts | 1 | 0.06 | 0.801 |
| | Treatment:particle type | 3 | 16.0 | **0.001** |
| | Treatment:extracts | 3 | 13.9 | **0.002** |
| | Particle type:extracts | 1 | 12.0 | **0.0005** |
| | Treatment:particle type:extracts | 3 | 16.2 | **0.0001** |
| ***ROS*** | Treatment | 3 | 8.4 | **0.037** |
| | Particle type | 1 | 2.0 | 0.147 |
| | Extracts | 1 | 1.0 | 0.315 |
| | Treatment:particle type | 3 | 2.7 | 0.428 |
| | Treatment:extracts | 3 | 0.4 | 0.928 |
| | Particle type:extracts | 1 | 1.2 | 0.260 |
| | Treatment:particle type:extracts | 3 | 1.5 | 0.669 |
| ***LPO*** | Treatment | 3 | 10.3 | **0.016** |
| | Particle type | 1 | 4.1 | **0.043** |
| | Extracts | 1 | 0.5 | 0.471 |
| | Treatment:particle type | 3 | 2.5 | 0.472 |
| | Treatment:extracts | 3 | 1.4 | 0.694 |
| | Particle type:extracts | 1 | 0.4 | 0.551 |
| | Treatment:particle type:extracts | 3 | 0.8 | 0.853 |

**Table SI 3:** Pairwise comparison between the separate polymers divided by treatment of the *D. magna*'s life history parameters obtained during the 10-day exposure. Bold values represent significance ($p < 0.05$) and a detectable effect ($D > 0.3$).

| *Contrasts* | *Cohen'D* | *t/z value* | *p value* |
|---|---|---|---|
| | ***Body length (day 6)*** | | |
| | ***MP*** | | |
| ***Cellulose-PBS*** | 0.211 | 0.747 | 1.000 |
| ***Cellulose-PDLLA*** | -0.165 | -0.586 | 1.000 |
| ***Cellulose-PET*** | -0.094 | -0.336 | 1.000 |
| ***PBS-PDLLA*** | **-0.376** | -1.333 | 1.000 |
| ***PBS-PET*** | **-0.306** | -1.083 | 1.000 |
| ***PDLLA-PET*** | 0.070 | 0.250 | 1.000 |
| | ***eMP*** | | |
| ***Cellulose-PBS*** | -0.072 | -0.256 | 1.000 |
| ***Cellulose-PDLLA*** | **-0.300** | -1.061 | 1.000 |
| ***Cellulose-PET*** | -0.078 | -0.276 | 1.000 |
| ***PBS-PDLLA*** | -0.227 | -0.805 | 1.000 |
| ***PBS-PET*** | -0.005 | -0.020 | 1.000 |
| ***PDLLA-PET*** | 0.222 | 0.785 | 1.000 |
| | ***Extracts*** | | |
| ***Cellulose-PBS*** | **0.533** | 1.885 | 0.060 |
| ***Cellulose-PDLLA*** | **3.500** | 12.377 | **<0.0001** |
| ***Cellulose-PET*** | **1.590** | 5.624 | **<0.0001** |
| ***PBS-PDLLA*** | **2.967** | 10.492 | **<0.0001** |
| ***PBS-PET*** | **1.057** | 3.739 | **0.0004** |
| ***PDLLA-PET*** | **-1.910** | -6.753 | **<0.0001** |
| | ***Num. of offspring*** | | |
| | ***MP*** | | |
| ***Cellulose-PBS*** | **0.618** | 3.093 | **0.009** |
| ***Cellulose-PDLLA*** | -0.181 | -1.678 | 0.186 |
| ***Cellulose-PET*** | 0.201 | 1.465 | 0.186 |
| ***PBS-PDLLA*** | **-0.799** | -4.136 | **0.0002** |
| ***PBS-PET*** | **-0.417** | -1.976 | 0.144 |
| ***PDLLA-PET*** | **0.382** | 2.999 | **0.010** |
| | ***eMP*** | | |
| ***Cellulose-PBS*** | -0.106 | -0.624 | 1.000 |
| ***Cellulose-PDLLA*** | -0.038 | -0.218 | 1.000 |
| ***Cellulose-PET*** | -0.136 | -0.813 | 1.000 |
| ***PBS-PDLLA*** | -0.067 | 0.408 | 1.000 |
| ***PBS-PET*** | -0.030 | -0.192 | 1.000 |
| ***PDLLA-PET*** | -0.097 | -0.599 | 1.000 |

| | | | |
|---|---|---|---|
| ***Extracts*** | | | |
| ***Cellulose-PBS*** | **0.533** | 2.853 | **0.017** |
| ***Cellulose-PDLLA*** | **3.500** | 4.316 | **0.0001** |
| ***Cellulose-PET*** | **1.590** | 3.467 | **0.003** |
| ***PBS-PDLLA*** | **2.967** | 1.825 | 0.203 |
| ***PBS-PET*** | **1.057** | 0.713 | 0.505 |
| ***PDLLA-PET*** | **-1.910** | -1.144 | 0.505 |
| ***ROS*** | | | |
| ***MP*** | | | |
| ***Cellulose-PBS*** | **0.690** | 1.028 | 1.000 |
| ***Cellulose-PDLLA*** | 0.203 | 0.322 | 1.000 |
| ***Cellulose-PET*** | **0.855** | 1.352 | 1.000 |
| ***PBS-PDLLA*** | **-0.486** | -0.725 | 1.000 |
| ***PBS-PET*** | 0.160 | 0.246 | 1.000 |
| ***PDLLA-PET*** | **-0.651** | -1.030 | 1.000 |
| ***eMP*** | | | |
| ***Cellulose-PBS*** | **1.257** | 2.016 | 0.240 |
| ***Cellulose-PDLLA*** | **0.520** | 0.822 | 0.828 |
| ***Cellulose-PET*** | **1.399** | 2.212 | 0.184 |
| ***PBS-PDLLA*** | **-0.755** | -1.194 | 0.711 |
| ***PBS-PET*** | 0.124 | 0.196 | 0.845 |
| ***PDLLA-PET*** | **-0.879** | -1.389 | 0.678 |
| ***Extracts*** | | | |
| ***Cellulose-PBS*** | **0.694** | 1.098 | 1.000 |
| ***Cellulose-PDLLA*** | -0.252 | -0.399 | 1.000 |
| ***Cellulose-PET*** | **0.350** | 0.533 | 1.000 |
| ***PBS-PDLLA*** | **-0.947** | -1.497 | 0.837 |
| ***PBS-PET*** | **-0.344** | -0.544 | 1.000 |
| ***PDLLA-PET*** | **-0.606** | -0.953 | 1.000 |
| ***LPO*** | | | |
| ***MP*** | | | |
| ***Cellulose-PBS*** | **-0.733** | -1.094 | 1.000 |
| ***Cellulose-PDLLA*** | -0.147 | -0.265 | 1.000 |
| ***Cellulose-PET*** | **-1.453** | -2.298 | 0.149 |
| ***PBS-PDLLA*** | **-0.566** | -1.072 | 1.000 |
| ***PBS-PET*** | **0.719** | 0.844 | 1.000 |
| ***PDLLA-PET*** | **1.286** | 2.033 | 0.231 |
| ***eMP*** | | | |
| ***Cellulose-PBS*** | **-0.985** | -1.262 | 0.846 |
| ***Cellulose-PDLLA*** | -0.026 | -0.554 | 1.000 |
| ***Cellulose-PET*** | **-0.795** | -1.954 | 0.331 |
| ***PBS-PDLLA*** | **-0.959** | -0.692 | 1.000 |
| ***PBS-PET*** | 0.190 | 0.708 | 1.000 |
| ***PDLLA-PET*** | **0.769** | 1.400 | 0.832 |
| ***Extracts*** | | | |
| ***Cellulose-PBS*** | **-0.733** | -1.559 | 0.744 |
| ***Cellulose-PDLLA*** | -0.167 | -0.042 | 1.000 |

| | | | |
|---|---|---|---|
| ***Cellulose-PET*** | **-1.453** | -1.257 | 0.853 |
| ***PBS-PDLLA*** | **0.566** | 1.517 | 0.744 |
| ***PBS-PET*** | **0.719** | 0.301 | 1.000 |
| ***PDLLA-PET*** | **1.285** | -1.215 | 0.853 |

**Table SI 4:** Pairwise comparison between the treatments divided by polymer of the *D. magna's* life history parameters obtained during the 10-day exposure. Bold values represent significance ($p < 0.05$) and a detectable effect ($D > 0.3$).

| *contrasts* | *Cohen'D* | *t/z value* | *p-value* |
|---|---|---|---|
| | ***Body length (day 6)*** | | |
| | ***Cellulose*** | | |
| ***Blank-MP*** | -0.146 | -0.516 | 1.000 |
| ***Blank-eMP*** | 0.095 | 0.337 | 1.000 |
| ***Blank-Extracts*** | -0.154 | -0.559 | 1.000 |
| ***eMP-Extracts*** | -0.253 | -0.896 | 1.000 |
| ***eMP-MP*** | -0.241 | -0.853 | 1.000 |
| ***Extracts-MP*** | 0.012 | 0.042 | 1.000 |
| | ***PBS*** | | |
| ***Blank-MP*** | 0.065 | 0.231 | 1.000 |
| ***Blank-eMP*** | 0.023 | 0.081 | 1.000 |
| ***Blank-Extracts*** | **0.375** | 1.326 | 1.000 |
| ***eMP-Extracts*** | **0.352** | 1.245 | 1.000 |
| ***eMP-MP*** | 0.042 | 0.150 | 1.000 |
| ***Extracts-MP*** | **-0.310** | -1.095 | 1.000 |
| | ***PDLLA*** | | |
| ***Blank-MP*** | **-0.312** | -1.102 | 0.813 |
| ***Blank-eMP*** | -0.205 | -0.724 | 0.939 |
| ***Blank-Extracts*** | **3.342** | 11.818 | **<0.0001** |
| ***eMP-Extracts*** | **3.548** | 12.542 | **<0.0001** |
| ***eMP-MP*** | -0.107 | -0.378 | 0.938 |
| ***Extracts-MP*** | **-3.310** | -12.920 | **<0.0001** |
| | ***PET*** | | |
| ***Blank-MP*** | -0.241 | -0.852 | 1.000 |
| ***Blank-eMP*** | 0.017 | 0.061 | 1.000 |
| ***Blank-Extracts*** | **1.433** | 5.065 | **<0.0001** |
| ***eMP-Extracts*** | **1.416** | 5.004 | **<0.0001** |
| ***eMP-MP*** | -0.256 | -0.913 | 1.000 |
| ***Extracts-MP*** | **-1.673** | -5.917 | **<0.0001** |
| | ***Num. of offspring*** | | |
| | ***Cellulose*** | | |
| ***Blank-MP*** | 0.203 | 1.901 | 0.115 |
| ***Blank-eMP*** | **0.540** | 3.756 | **0.0009** |
| ***Blank-Extracts*** | -0.135 | -1.565 | 0.117 |
| ***eMP-Extracts*** | **-0.675** | -4.837 | **<0.0001** |

| | | | |
|---|---|---|---|
| ***eMP-MP*** | **-0.337** | -2.199 | 0.083 |
| ***Extracts-MP*** | **0.338** | 3.343 | **0.003** |
| | ***PBS*** | | |
| ***Blank-MP*** | **0.821** | 4.263 | **0.001** |
| ***Blank-eMP*** | **0.434** | 3.338 | **0.003** |
| ***Blank-Extracts*** | 0.138 | 1.358 | 0.175 |
| ***eMP-Extracts*** | -0.256 | -2.165 | 0.091 |
| ***eMP-MP*** | **0.387** | 1.816 | 0.139 |
| ***Extracts-MP*** | **0.682** | 3.464 | **0.003** |
| | ***PDLLA*** | | |
| ***Blank-MP*** | 0.022 | 0.232 | 0.869 |
| ***Blank-eMP*** | **0.502** | 3.621 | **0.002** |
| ***Blank-Extracts*** | **0.376** | 3.048 | **0.009** |
| ***eMP-Extracts*** | -0.126 | -0.781 | 0.869 |
| ***eMP-MP*** | **-0.480** | -3.441 | **0.003** |
| ***Extracts-MP*** | **-0.354** | -2.849 | **0.013** |
| | ***PET*** | | |
| ***Blank-MP*** | **0.404** | 3.194 | **0.008** |
| ***Blank-eMP*** | **0.404** | 2.040 | **0.008** |
| ***Blank-Extracts*** | 0.221 | 3.194 | 0.165 |
| ***eMP-Extracts*** | -0.183 | -1.322 | 0.558 |
| ***eMP-MP*** | 0.000 | 0.000 | 1.000 |
| ***Extracts-MP*** | 0.183 | 1.322 | 0.558 |
| | ***ROS*** | | |
| | ***Cellulose*** | | |
| ***Blank-MP*** | 0.071 | 0.105 | 1.000 |
| ***Blank-eMP*** | **-0.776** | -1.146 | 1.000 |
| ***Blank-Extracts*** | -0.233 | -0.344 | 1.000 |
| ***eMP-Extracts*** | **0.543** | 0.802 | 1.000 |
| ***eMP-MP*** | **0.847** | 1.251 | 1.000 |
| ***Extracts-MP*** | **0.304** | 0.449 | 1.000 |
| | ***PBS*** | | |
| ***Blank-MP*** | **0.867** | 1.127 | 1.000 |
| ***Blank-eMP*** | **0.589** | 0.870 | 1.000 |
| ***Blank-Extracts*** | **0.510** | 0.754 | 1.000 |
| ***eMP-Extracts*** | -0.078 | -0.116 | 1.000 |
| ***eMP-MP*** | 0.277 | 0.306 | 1.000 |
| ***Extracts-MP*** | **0.356** | 0.416 | 1.000 |
| | ***PDLLA*** | | |
| ***Blank-MP*** | 0.288 | 0.426 | 1.000 |
| ***Blank-eMP*** | -0.219 | -0.324 | 1.000 |
| ***Blank-Extracts*** | **-0.503** | -0.743 | 1.000 |
| ***eMP-Extracts*** | -0.284 | -0.419 | 1.000 |

| | | | |
|---|---|---|---|
| ***eMP-MP*** | **0.508** | 0.750 | 1.000 |
| ***Extracts-MP*** | **0.792** | 1.169 | 1.000 |
| ***PET*** | | | |
| ***Blank-MP*** | **0.987** | 1.456 | 0.902 |
| ***Blank-eMP*** | **0.722** | 1.066 | 1.000 |
| ***Blank-Extracts*** | 0.142 | 0.210 | 1.000 |
| ***eMP-Extracts*** | **-0.580** | -0.856 | 1.000 |
| ***eMP-MP*** | 0.265 | 0.391 | 1.000 |
| ***Extracts-MP*** | **0.845** | 1.247 | 0.934 |
| ***LPO*** | | | |
| ***Cellulose*** | | | |
| ***Blank-MP*** | -0.018 | -0.028 | 1.000 |
| ***Blank-eMP*** | 0.279 | 0.441 | 1.000 |
| ***Blank-Extracts*** | **0.418** | 0.662 | 1.000 |
| ***eMP-Extracts*** | 0.139 | 0.221 | 1.000 |
| ***eMP-MP*** | -0.296 | -0.469 | 1.000 |
| ***Extracts-MP*** | **-0.437** | -0.691 | 1.000 |
| ***PBS*** | | | |
| ***Blank-MP*** | **-0.751** | -1.121 | 1.000 |
| ***Blank-eMP*** | **-0.519** | -0.821 | 1.000 |
| ***Blank-Extracts*** | **-0.567** | -0.897 | 1.000 |
| ***eMP-Extracts*** | -0.048 | -0.075 | 1.000 |
| ***eMP-MP*** | -0.232 | -0.346 | 1.000 |
| ***Extracts-MP*** | -0.185 | -0.275 | 1.000 |
| ***PDLLA*** | | | |
| ***Blank-MP*** | -0.185 | -0.293 | 1.000 |
| ***Blank-eMP*** | -0.071 | -0.113 | 1.000 |
| ***Blank-Extracts*** | **0.392** | 0.620 | 1.000 |
| ***eMP-Extracts*** | **0.463** | 0.734 | 1.000 |
| ***eMP-MP*** | -0.113 | -0.180 | 1.000 |
| ***Extracts-MP*** | **-0.577** | -0.913 | 1.000 |
| ***PET*** | | | |
| ***Blank-MP*** | **-1.471** | -2.326 | 0.139 |
| ***Blank-eMP*** | **-0.957** | -1.513 | 0.541 |
| ***Blank-Extracts*** | **-0.376** | -0.595 | 1.000 |
| ***eMP-Extracts*** | **0.580** | 0.918 | 1.000 |
| ***eMP-MP*** | **-0.514** | -0.813 | 1.000 |
| ***Extracts-MP*** | **-1.095** | -1.713 | 0.442 |

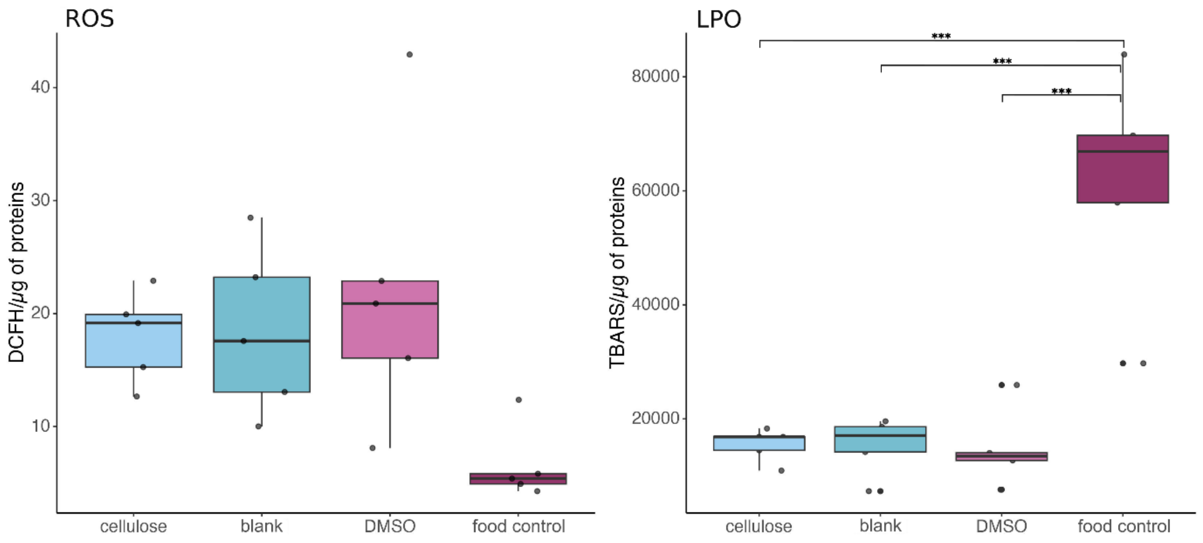


***Figure SI 11***: *Levels of ROS and LPO (N=5) recorded for the control treatments, grouped by polymer and treatment type (MP, eMP, and extracts). Horizontal lines indicate the median for each treatment, while the boxes represent the interquartile range. Individual data points correspond to values obtained for each replicate. Asterisks indicate significance against the blank control (*** = $p \leq 0.001$).*

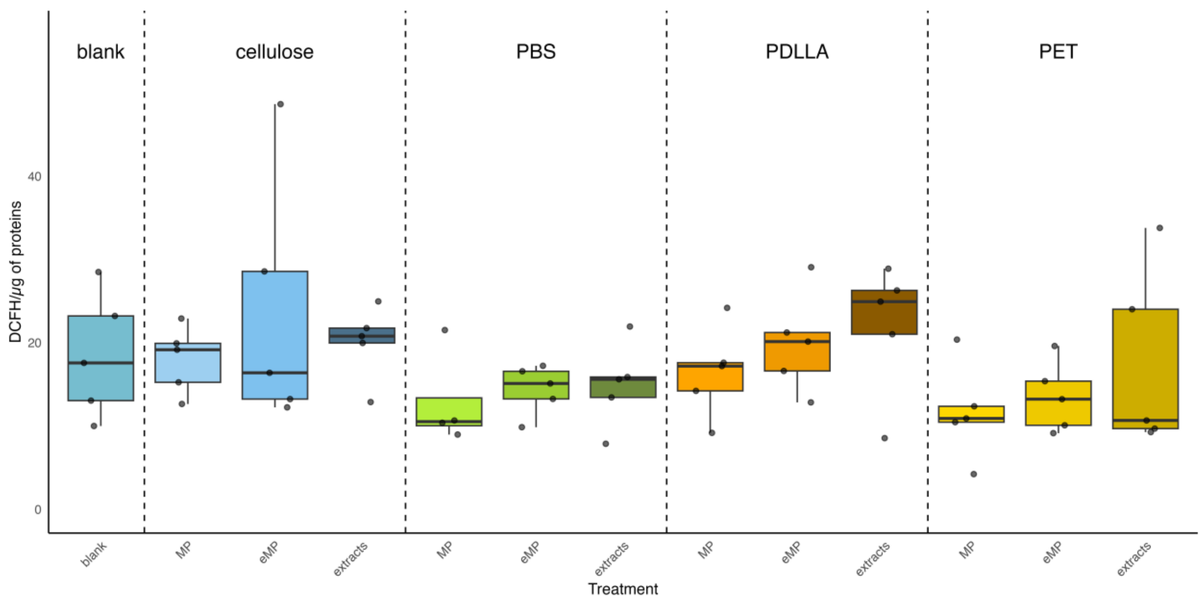

***Figure SI 12****: Levels of ROS measured at the end of the exposure expressed in DCFH/µg of proteins grouped by polymer and treatment type (MP, eMP, and extracts). N=5. Horizontal lines indicate the median for each treatment, while the boxes represent the interquartile range. Individual data points correspond to values obtained for each replicate.*

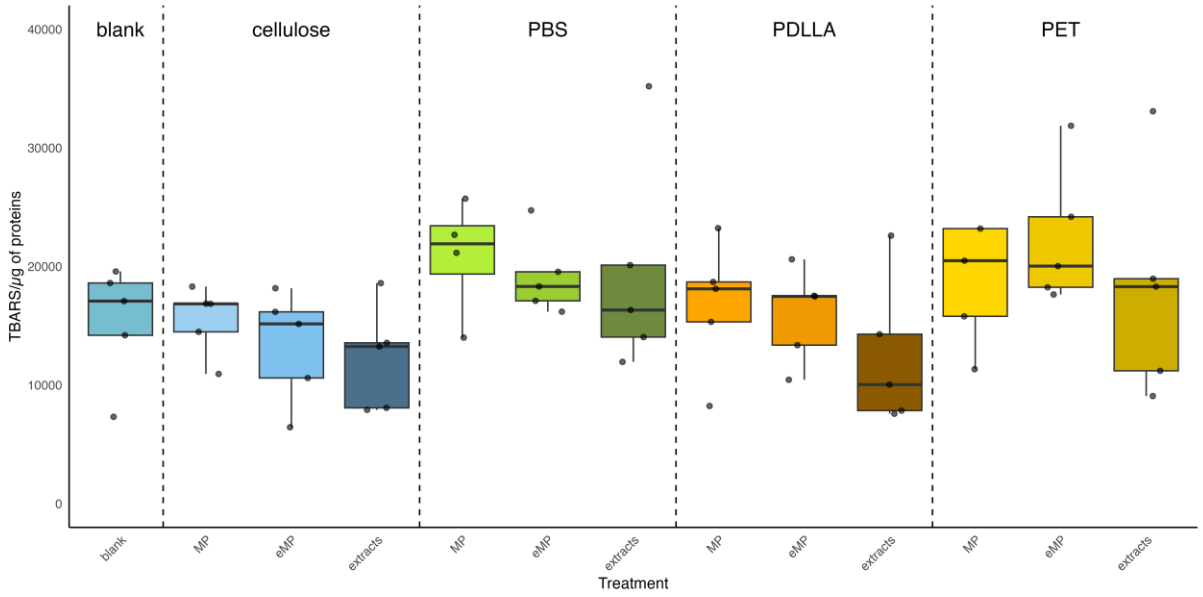


***Figure SI 13:*** *Levels of LPO measured at the end of the exposure expressed in TBARS/µg of proteins grouped by polymer and treatment type (MP, eMP, and extracts). N=5. Horizontal lines indicate the median for each treatment, while the boxes represent the interquartile range. Individual data points correspond to values obtained for each replicate.*

### Discussion on GC-MS results

GC-MS analyses of the extracts showed several plastic chemicals. We were able to confirm the identity of four of them when the calculated Kovats indices matched those reported in the literature. Exposure to mixtures of plastic chemicals, such as those found in our extracts, could significantly affect organisms, since these chemicals may interact in antagonistic, additive, or synergistic ways, even if they are not individually identified as potentially harmful. This is due to the complex interactions among the chemicals present in the mixtures (Silva et al. 2022), which make it difficult to predict outcomes solely from their chemical composition. These complex interactions should be better assessed in future investigations. In total, 19 chemicals were detected inside the employed extracts. PBS contained the most chemicals, with 10, followed by PET and PDLLA (4 each), and cellulose, which contained only 1.

PDLLA had the same number of chemicals as PET, and the BB-MP PBS had the highest number of associated chemicals in our study. This may be because, particularly in BB-plastics, mixtures of several chemicals are often used to achieve properties comparable to those of PB-plastics (Savva et al. 2023).

The employed extraction protocol should ensure that most extractable chemicals are obtained from the original material using an organic solvent (i.e., methanol; Zimmermann et al. 2020). Therefore, the obtained peaks should be representative of the chemicals incorporated into pre-production pellets used in this study.

One of these chemicals was Thiodiglycol, which was detected in both PET and PBS. This compound is commonly used in plastic manufacturing, particularly to manipulate the microstructures of polyesters (Fuoco and Finne-Wistrand 2020), such as PET and PBS. It is considered to exhibit low toxicity, with a reported $EC_{50}$ on *D. magna* above 500 mg/L (OECD 2004). In PET, we also identified docosane. This compound is an n-alkane commonly detected in plastic debris, and also reported in PlastChem (https://plastchem-project.org/, Wagner et al. 2025). Like other alkanes, docosane is used either as an external lubricant, to help polymers slide over one another, or as a solvent and starting compound for diverse additives and polymers, particularly during olefin polymerization (Espinosa, Esteban, and Cuesta 2016). This compound is generally regarded as of low acute hazard (ECHA, 2018). The GC-MS analyses of PBS extracts also highlighted the presence of dimethyl ester of butanedioic acid. This compound is used as a monomer in the synthesis of polyesters (Miller, Nguyen, and Short

2021; Nikolic and Djonlagic 2001), is also reported in the PlastChem repository (https://plastchem-project.org/, Wagner et al. 2025), and is generally considered to be of low toxicity. According to ECHA (2021), no effect was observed in a 48-hour acute toxicity test conducted in accordance with OECD Guideline 202 (2004). Nevertheless, it is classified as an eye irritant and potentially harmful to the upper respiratory tract in humans (NCBI).

In PDLLA, four peaks were identified, but the identity of only one of them, 3-methyl-2(5H)-furanone, could be confirmed. While this compound is commonly employed in the flavor and fragrance industry (Effenberger et al. 2019), there is no evidence that it is used in the process of plastic manufacturing. Data on its potential effects are very limited, and no data are available for aquatic organisms, but it is reported to be a skin, eye, and respiratory tract irritant in humans (Sigma-Aldrich, 2026).

One peak was also found in the cellulose extracts. However, we were unable to identify this compound with certainty. Nevertheless, the cellulose extract treatment had no adverse effects on the exposed daphnids.

**Limitations and recommendations**

This study revealed some limitations and recommendations for future studies: (i) The exposure duration was limited to only 10 days, and thus, only the first brood of the tested animals was assessed. Future investigations should extend exposure periods in order to capture potential longer-term or multigenerational effects. Standard chronic toxicity testing protocols (OECD No. 211), modified to incorporate more environmentally relevant scenarios and extended to 21 days or longer (e.g., multigenerational), may provide a more comprehensive assessment. (ii) Reduced food availability led to smaller brood sizes, substantially decreasing the number of produced neonates. Although this reduced the overall sample size, it represents a more environmentally relevant scenario, potentially better reflecting the effects of plastic exposure under natural conditions, where food limitation can co-occur with contaminant stressors. (iii) As a mechanistic study, this study used a relatively high concentration that exceeded those found in the environment (Gupta et al. 2022, Moses et al. 2023, Langenfeld et al. 2024), limiting the extent to which the results could be applied to environmentally relevant scenarios. (iv) In the current study, plastic chemicals exerted the most prominent effects. However, the chemicals detected by GC-MS analysis alone were likely not responsible for the observed toxicity. The applied GC-MS protocol enabled only a qualitative assessment of the chemicals present in the extracts. Furthermore, this technique has limited sensitivity to non-volatile compounds (Beale et al. 2018), indicating that not all chemicals present in the extracts could be measured. Volatility of analytes can be increased by derivatization, but this is only feasible for target analyses, as such reactions are inherently selective and introduce compound-class-dependent biases. This limits their applicability in untargeted analysis, making them more suitable for targeted approaches (Moldoveanu and David 2018). (v) Extractable chemicals generally represent a worst-case scenario compared to leachates, as extraction protocols can recover a broader range of compounds at higher concentrations, whereas leaching of plastic chemicals requires longer incubations but probably better simulates realistic release processes in the environment. Nonetheless, while leachates might be more informative for an ecotoxicological risk assessment, extractable chemicals provide a comprehensive overview of the compounds associated with MP (Bridson et al. 2023), giving precise indications of their toxic potential.

Organisation for Economic Co-operation and Development (OECD). 2004. “Test No. 202: *Daphnia* sp. acute immobilisation test.” OECD Publishing. 10.1787/9789264069947-en.1

Organisation for Economic Co-operation and Development (OECD) and United Nations Environment Programme (UNEP). 2004. SIDS Initial Assessment Report for Thiodiglycol (CAS No. 111-48-8). OECD HPV Chemicals Database. https://hpvchemicals.oecd.org/ui/handler.axd?id=fa33e917-fe0c-4d58-8ffd-2ba605d502b31

Savva, K., Borrell, X., Moreno, T., Pérez-Pomeda, I., Barata, C., Llorca, M., and Farré, M. 2023. “Cytotoxicity Assessment and Suspected Screening of PLASTIC ADDITIVES in Bioplastics of Single-Use Household Items.” *Chemosphere* 313: 137494. 10.1016/j.chemosphere.2022.137494.1

Sigma-Aldrich. 2026. *Safety Data Sheet: 3-Methyl-2(5H)-furanone (Product No. 393509).* Merck KGaA.

Silva, A. R. R., Gonçalves, S. F., Pavlaki, M. D., Morgado, R. G., Soares, A. M. V. M., and Loureiro, S. 2022. “Mixture Toxicity Prediction of Substances from Different Origin Sources in *Daphnia magna*.” *Chemosphere* 292: 133432. 10.1016/j.chemosphere.2021.133432.

Wagner, M., Monclús, L., Arp, H. P. H., Groh, K. J., Løseth, M. E., Muncke, J., Wang, Z., Wolf, R., and Zimmermann, L. 2025. “State of the science on plastic chemicals - Identifying and addressing chemicals and polymers of concern (1.01).” Zenodo. 10.5281/zenodo.17208791.

Zimmermann, L., Göttlich, S., Oehlmann, J., Wagner, M., and Völker, C. 2020. “What Are the Drivers of Microplastic Toxicity? Comparing the Toxicity of Plastic Chemicals and Particles to *Daphnia magna*.” *Environmental Pollution* 267: 115392. 10.1016/j.envpol.2020.115392.
